# Time resolved quantitative phosphoproteomics reveals distinct patterns of SHP2 dependence in EGFR signaling

**DOI:** 10.1101/598664

**Authors:** Vidyasiri Vemulapalli, Lily Chylek, Alison Erickson, Jonathan LaRochelle, Kartik Subramanian, Morvarid Mohseni, Matthew LaMarche, Michael G. Acker, Peter K. Sorger, Steven P. Gygi, Stephen C. Blacklow

## Abstract

SHP2 is a protein tyrosine phosphatase that normally potentiates intracellular signaling by growth factors, antigen receptors, and some cytokines; it is frequently mutated in childhood leukemias and other cancers. Here, we examine the role of SHP2 in the responses of breast cancer cells to EGF by monitoring phosphoproteome dynamics when SHP2 is allosterically inhibited by the small molecule SHP099. Data on phosphotyrosine abundance at more than 400 tyrosine residues reveals six distinct response signatures following SHP099 treatment and washout. These include putative substrate sites with increased phosphotyrosine abundance at early or late timepoints, and another class of sites that shows reduced phosphotyrosine abundance when SHP099 is present. Sites of decreased phospho-abundance are enriched on proteins with two nearby phosphotyrosine residues, which can be directly protected from dephosphorylation by the paired SH2 domains of SHP2 itself. These findings highlight how analysis of phosphoproteomic dynamics can provide insight into transmembrane signaling responses.

## Introduction

SHP2 is a non-receptor tyrosine phosphatase that is essential for mammalian development (Saxton et al., 1997). In humans, germline mutations of *PTPN11* cause the developmental disorders Noonan and LEOPARD syndromes (Tartaglia & Gelb, 2005). Somatic *PTPN11* activating mutations are also found frequently in Juvenile Myelomonocytic Leukemia (JMML) and, to a lesser extent, in solid tumors (Bentires-Alj et al., 2004).

Numerous studies have shown that SHP2 acts as a positive effector of receptor tyrosine kinase (RTK) signaling (Bennett, Tang, Sugimoto, Walsh, & Neel, 1994; Easton, Royer, & Middlemas, 2006; Tang, Freeman, O’Reilly, Neel, & Sokol, 1995). SHP2 facilitates the full induction of Ras-dependent extracellular signal-regulated kinase (ERK) proteins following stimulation of cells with epidermal growth factor (EGF) or other receptor tyrosine kinase ligands. SHP2 also serves as a positive regulator of numerous other signaling systems, including cytokine (Xu & Qu, 2008), programmed cell death (PD-1) (Yokosuka et al., 2012), and immune checkpoint pathways (Gavrieli, Watanabe, Loftin, Murphy, & Murphy, 2003).

The SHP2 protein consists of two SH2 domains (N-SH2, C-SH2), followed by a phosphatase (PTP) domain, with a natively disordered C-terminal tail that contains tyrosine residues known to become phosphorylated (Bennett et al., 1994; Keegan & Cooper, 1996). X-ray structures of the SHP2 protein core, encompassing the two SH2 domains and the PTP domain, show that the wild-type protein normally adopts an autoinhibited conformation in which the N-SH2 domain occludes the active site of the PTP domain (Hof, Pluskey, Dhe-Paganon, Eck, & Shoelson, 1998). Activation of the enzyme is thought to require disengagement of the N-SH2 domain from the PTP domain and subsequent recruitment of SHP2 to phosphotyrosine docking sites on substrate proteins; recruitment can involve the N-SH2 domain, C-SH2 domain, or both domains. Most oncogenic mutations of SHP2 lie at the interface between the N-SH2 and PTP domains, thereby increasing the propensity of the enzyme to adopt an open, active conformation (LaRochelle et al., 2018; LaRochelle et al., 2016).

Numerous studies in a range of cell types have identified phosphotyrosine (pY) residues on adaptor proteins (*e.g.* GAB1 and GAB2 (Arnaud et al., 2004; Cunnick, Mei, Doupnik, & Wu, 2001)) or on RTKs themselves (*e.g.* PDGFβR (Ronnstrand et al., 1999)) thought to recruit SHP2 to sites of active signaling following RTK activation (such sites include multiprotein complexes formed on RTK intracellular tails and IRS-1 adapters). Genetic and mutational studies have implicated specific pY residues on RTKs as putative SHP2 substrates (Agazie & Hayman, 2003; Bunda et al., 2015; Klinghoffer & Kazlauskas, 1995). Because a potent and highly selective inhibitor of SHP2 was previously unavailable, however, previous studies investigating the influence of SHP2 on the response to EGF were primarily limited to candidate-based approaches for identification of substrates and recruitment motifs.

Here, we use the recently reported allosteric inhibitor SHP099 (Garcia Fortanet et al., 2016) to study the role of SHP2 on phosphoprotein dynamics following EGF stimulation of a breast cancer cell line carrying an EGFR amplification. Global analysis of time-resolved changes in phosphotyrosine abundance at over 400 tyrosine residues reveals several distinguishable response signatures following SHP099 exposure and washout. Putative substrate sites fall into two classes, with increased phosphotyrosine abundance at early and/or late timepoints, respectively. Yet another class, which contains the largest number of dynamic pY sites, exhibits decreased phosphotyrosine abundance. These latter sites are enriched on proteins with two nearby phosphotyrosine residues that may be directly protected from dephosphorylation by the SH2 domains of SHP2. These data emphasize the two competing biochemical activities of SHP2 - dephosphorylation and phosphosite protection – and identify specific sites relevant to the activity of SHP099 and similar molecules as therapeutics.

## Results

### Dynamic regulation of the EGF-responsive phosphoproteome by SHP2

We investigated the role of SHP2 in the responsiveness to EGF by using the EGFR-amplified cell line MDA-MB-468, derived from a patient with triple negative breast cancer (TNBC). To identify timepoints for in-depth proteomic analysis, we monitored the influence of SHP2 on the response to EGF stimulation using ERK1/2 phosphorylation as a readout. Cell extracts were prepared at a series of timepoints from three treatment conditions: i) following pretreatment with DMSO and then stimulation with 10 nM EGF, ii) following pretreatment with the SHP2 inhibitor SHP099 for 2 hours and then stimulation with EGF, and iii) following pretreatment with SHP099 and then stimulation with EGF for 10 min, after which drug was washed out and medium containing EGF replenished. Immunoblot analysis revealed the expected effect of EGF stimulation: a dramatic increase in phospho-ERK1/2 (p-ERK1/2) levels followed by a decline toward basal levels by 30 min. The induction of p-ERK1/2 was greatly attenuated by pretreatment of cells with 10 μM SHP099. SHP099 washout in the continued presence of EGF (condition iii) revealed p-ERK1/2 induction with a kinetic profile similar to that of EGF stimulation in the absence of drug (condition i), showing that SHP099 inhibition of SHP2 is rapidly reversible in MDA-MB-468 cells **(Figure 1A)**.

**Figure 1.**
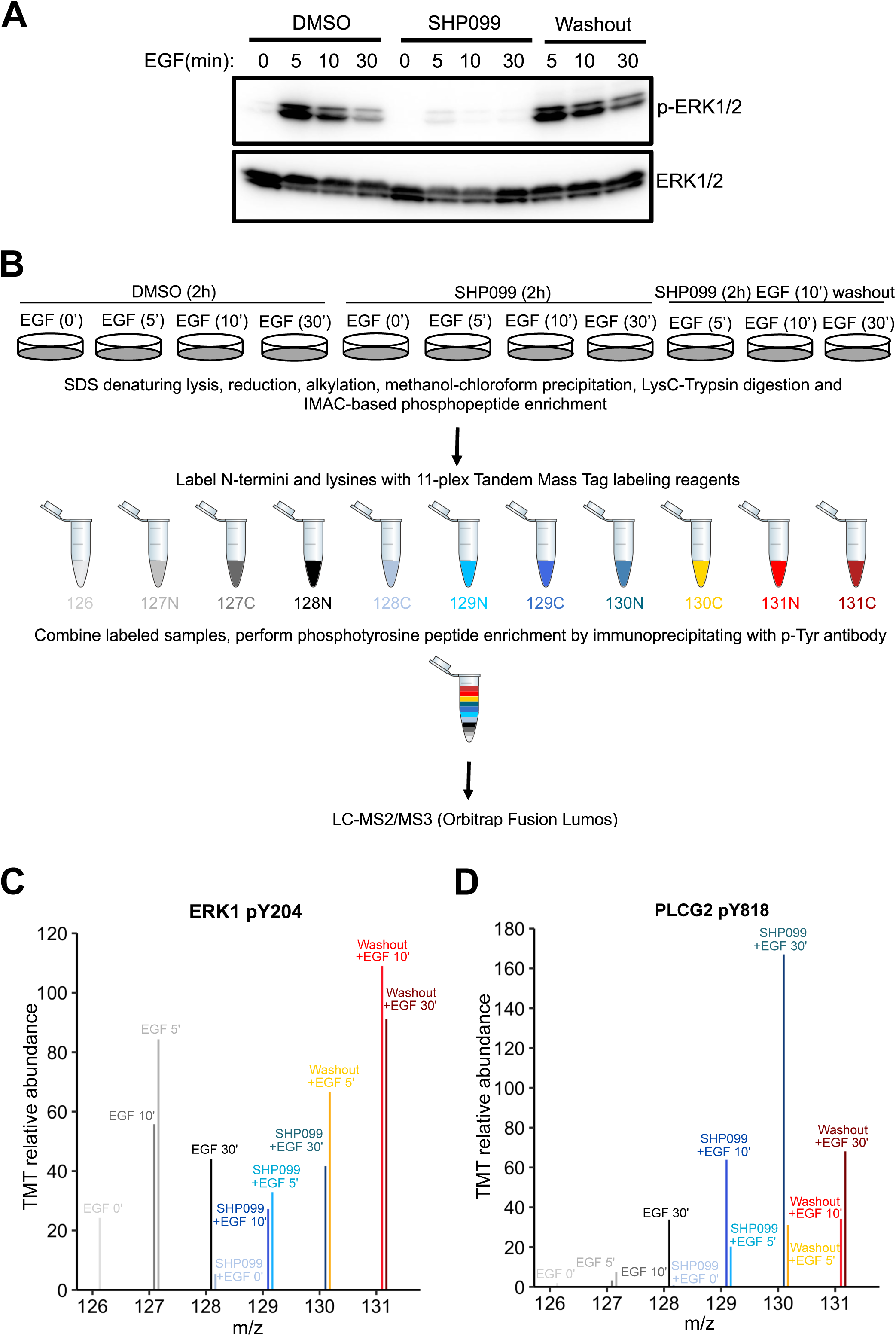
Phosphoproteomic studies in MDA-MB-468 cells. (A) Western blot showing the phosphoERK1/2 abundance after EGF stimulation alone, EGF stimulation in the presence of SHP099, and EGF stimulation in the presence of SHP099, followed by drug washout 10 minutes after EGF stimulation. (B) Schematic illustration of treatment conditions and mass spectrometry workflow. (C, D) TMT relative abundance plotted as a function of peptide m/z values from MS3 spectra, showing quantification of ERK1 (C) and PLCG2 (D) phosphopeptides.

To obtain an in-depth view of the effects of SHP2 inhibition on EGFR signaling we performed quantitative phosphoproteomics, monitoring dynamic changes in phosphotyrosine abundance as a function of time under DMSO, SHP099, and washout conditions **(Figure 1B**). Tryptic peptides from DMSO, SHP099, and SHP099-washout groups were enriched for phosphopeptides using Immobilized Metal Affinity Chromatography (IMAC) prior to labeling with 11-plex isobaric tandem mass tags (TMT). Tyrosine phosphorylated (pY) peptides were then immunoprecitiated from the TMT-labelled, pooled samples using an anti-phosphotyrosine antibody, and the recovered pY-containing peptides were analyzed by LC-MS3 mass spectrometry (**Table S1 and Figures 1C, D**). This experiment yielded relative quantification for several hundred phosphotyrosine-containing peptides (496 and 748 localized and quantified sites in the first and second experimental replicates, respectively; **Table S1**). Results for the dynamics of pY204 of ERK1 **(Figure 1C)** and pY187 of ERK2 **(Table S1)** modification obtained by mass spectrometry experiments matched the results of Western blotting described above **(Figure 1A)**, confirming that key phosphotyrosine marks associated with EGF-induced signaling events were accurately determined in the TMT experiment.

The phosphoproteomic data revealed a spectrum of dependencies on EGF and SHP2 **(Figure S1A)**. These were classified on the basis of both direction of change (increase or decrease) and timing. EGF dependencies were established based on responses to EGF addition in the absence of drug, and were classified into five categories: fast, medium, or slow increases, neutral (no significant change), or decrease. The dependence of each phosphosite on SHP2 activity was established by assessing how phosphorylation levels were altered when SHP2 was inhibited: negative (more phosphorylation with inhibition; Figure S1A, top two rows), positive (less phosphorylation with inhibition; Figure S1A, bottom two rows), or neutral (Figure S1A, middle row). Positive and negative dependencies were further subdivided based on whether the change was present before EGF stimulation (Figure S1A, first and fourth rows) or only after (Figure S1A, second and fifth rows).

As anticipated, SHP2 inhibition resulted in increased abundance of phosphotyrosine marks at a number of sites **(Figure S1A, top two rows)**, the expected effect of blocking a tyrosine phosphatase. Remarkably, however, we found an even larger family of phosphotyrosine sites that showed a decreased abundance when SHP2 is inhibited even at early time points **(Figure S1A, bottom two rows)**, indicating that SHP2 inhibition had direct and/or indirect activities other than direct dephosphorylation of substrates. Additionally, SHP2 exhibited EGFR-independent regulation of a number of pY sites: a subset of sites was regulated (either positively or negatively) by SHP2 prior to EGF addition **(Figure S1A, first and fourth rows)**, showing that SHP2 had a role in shaping the basal signaling state of these cells. Motif analysis of phosphosites associated with different classes of SHP2 responsiveness did not reveal enrichment for specific motifs (residues flanking the modified tyrosine residue**; Figure S1B)**. Remarkably, sites from different regulatory classes could even occur together in the same protein (see below), highlighting the importance of considering specific sites of phosphorylation when evaluating the impact of a phosphatase on a protein substrate.

Hierarchical clustering was performed to identify pY sites with similar kinetics in DMSO, SHP099, and washout conditions **(Figure 2).** Among the six kinetic profiles that emerged **(Figure S2)**, we highlight three clusters of sites implicated in EGFR signaling and that display distinct responses to the drug **(Figure 2A)**. The first class of responses, which includes the regulatory subunits of PI3K, Syntaxin 4, and the adapter proteins CRK and GRB2, among other proteins, shows quantitatively increased tyrosine phosphorylation in the presence of SHP099 that rapidly disappears upon drug washout (**Figure 2A,B**). The second class of sites accumulates pY marks slowly under SHP099 inhibition (at 30 min after EGF treatment), and not when SHP099 is omitted or washed out (**Figure 2A,C**). This pattern, observed for phosphotyrosines on CBL E3 ligase, ANXA2 and RAB10 endocytic proteins, hnRNPs, and PLCγ proteins, also matches the response predicted for a SHP2 substrate, but with a time delay suggesting that the action of SHP2 on these proteins might require an intervening event, such as relocalization (for example, after endocytosis of active EGFR signaling complexes), or alternatively, that the effect is indirect. Among the members of these two classes, only two proteins have previously been reported to be dephosphorylated by SHP2 (ARHGAP35 pY1105 and GRB2 pY209) (Bregeon, Loirand, Pacaud, & Rolli-Derkinderen, 2009; Chardin et al., 1993). The large number of potential substrates newly identified in this work strongly suggests that SHP2 catalyzes dephosphorylation of many different proteins on different time scales.

**Figure 2.**
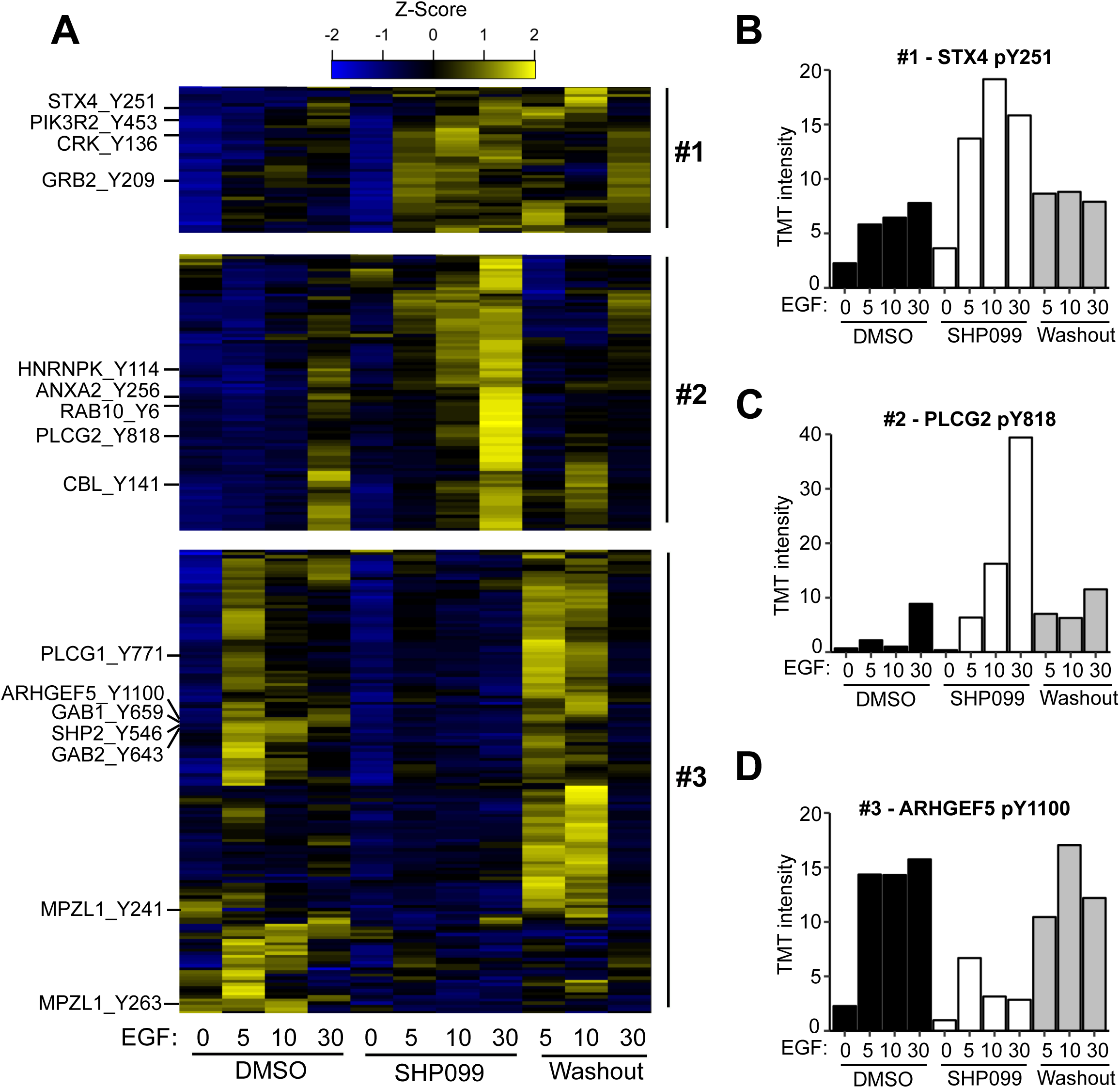
Quantitative phosphoproteomics reveals distinct dynamic responses to SHP2 inhibition. (A) Heatmap showing three classes of dynamic response in which inhibition of SHP2 modulates the effect of EGF stimulation on pY abundance. Specific examples from each cluster are indicated to the left of the heatmap. (B) Plot of pY abundance as a function of treatment condition for Y251 of STX4, an example of an early substrate-like response pattern to SHP2 inhibition. (C) Plot of pY abundance as a function of treatment condition for Y818 of PLCG2, an example of a late substrate-like response pattern to SHP2 inhibition. (D) Plot of pY abundance as a function of treatment condition for Y1100 of ARHGEF5, an example of a site where the abundance of the mark decreases when SHP2 is inhibited, and rebounds after compound washout.

The third pattern of response observed is one in which accumulation of phosphotyrosine in response to EGF *depends* on release of SHP2 from inhibition by SHP099. Examples of sites that fall into this category include Y659 of GAB1, Y643 of GAB2, as well as Y546 and Y584 of SHP2 itself (**Figure 2A,D**).

Functional classification of EGF-responsive pY sites (by GSEA and Reactome) differentially regulated by SHP099 **(Figure 3A-C)** reveals enrichment of six major cellular processes **(Figure 3D)**. As expected, proteins implicated in MAPK and PI3K signaling, including EGFR and SHP2 itself, contain numerous EGF-responsive phosphosites differentially regulated by SHP099 treatment. The largest functional group includes proteins implicated in adhesion and migration, including tight junction proteins, catenins, Rho-GEF, and Rho-GAP proteins. These findings, along with data on the abundance of cytoskeletal proteins differentially affected by SHP099 **(Figure 3D)**, are consistent with earlier results suggesting that SHP2 promotes migration in MDA-MB-468 cells by regulating EGF-induced lamellipodia persistence (Hartman, Schaller, & Agazie, 2013). Proteins implicated in transcription comprise another functional class with a number of differentially regulated phosphosites (**Figure 3D**). Two additional functional categories enriched in the analysis comprise endocytosis and mRNA processing, neither of which have previously been linked to SHP2 activity. Most of the sites in these two categories display a substrate-like response with SHP099 treatment. In addition, SHP099 also affected phosphorylation of CBL and CBLB, E3 ligases that ubiquitinate EGFR and facilitate its recruitment to clathrin-coated pits for endocytosis. Together, these results highlight the diversity of influence of SHP2, both in the function of proteins that it affects, and the manner in which it affects their phosphorylation patterns.

**Figure 3.**
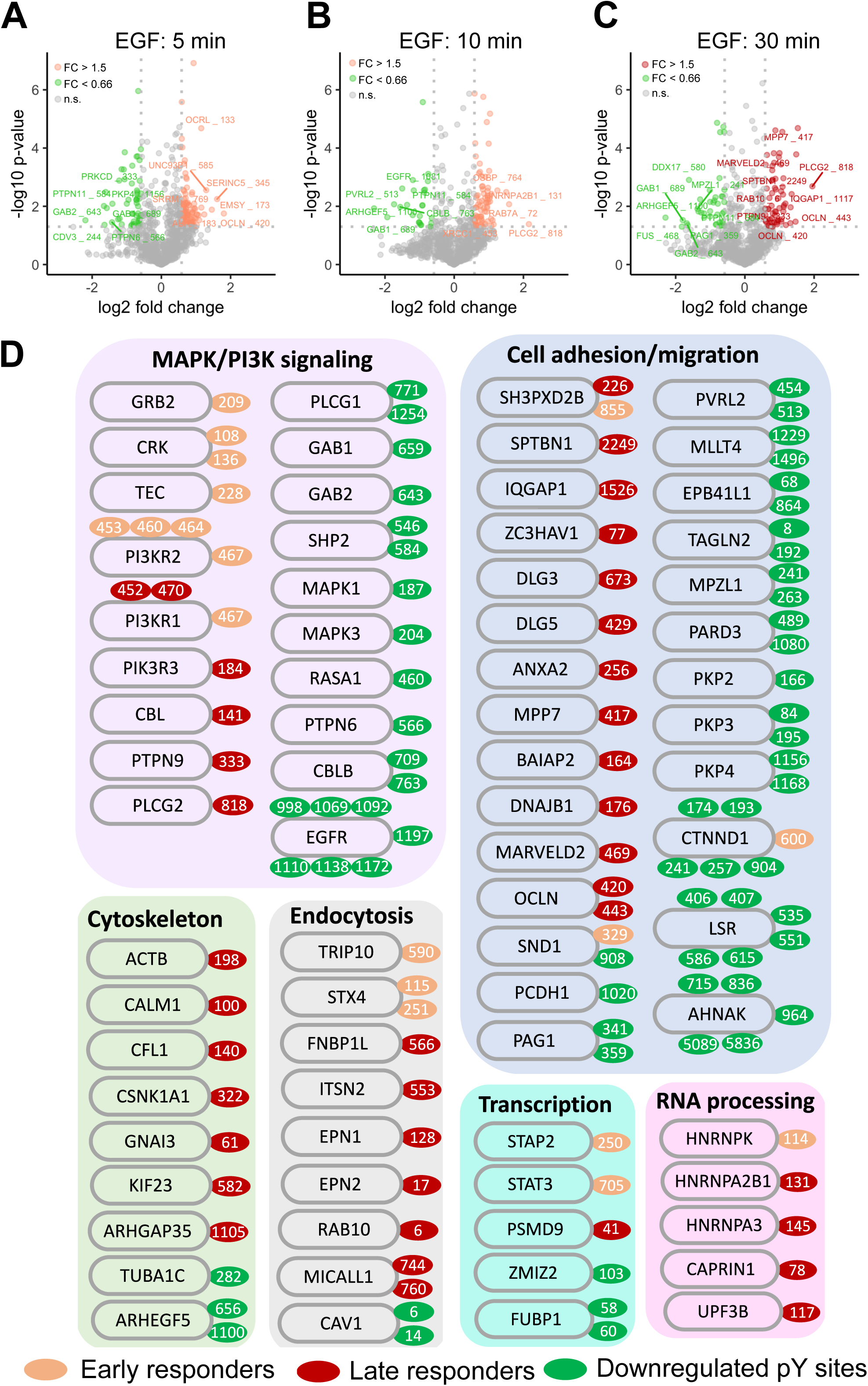
Classification of SHP099-responsive phosphotyrosine sites based on EGF temporal dynamics and function. (A-C) Semi-log volcano plots of p value as a function of pY fold-change after EGF stimulation for 5 min (A), 10 min (B), and 30 min (C). Sites exhibiting biologically significant increases (p < 0.05 and increase > 1.5 fold) are colored salmon (A,B) or red (C). Sites exhibiting statistically biologically significant decreases (p < 0.05 and decrease > 1.5 fold) are green. (D) Functional classification of proteins whose pY sites display statistically significant changes in response to SHP099 treatment. Classes were assigned using Gene Ontology categories and literature curation.

### Allosteric inhibition of SHP2 results in reduced pY abundance at its interaction motifs

SHP099 allosterically stabilizes the autoinhibited conformation of SHP2 (Chen et al., 2016), thereby not only inhibiting the catalytic activity of the enzyme, but also suppressing binding of its two SH2 domains to phosphotyrosine-containing motifs. Because the third pattern of response shows accumulation of phosphotyrosine after SHP2 is released from inhibition, we tested whether SHP2 directly protects sites in this class from dephosphorylation and whether this protection is lost upon SHP099 binding. We focused on Y659 of GAB1 and Y643 of GAB2, because both sites show dramatic reductions in pY abundance upon SHP099 treatment and recover their pY marks upon compound washout **(Figure 4),** and because prior work indicates that the SH2 domains of SHP2 can bind to the bisphosphotyrosine-containing motifs of GAB1 (pY627/pY659) and GAB2 (pY614/pY643) to relieve SHP2 autoinhibition (Arnaud et al., 2004; Cunnick et al., 2001).

**Figure 4.**
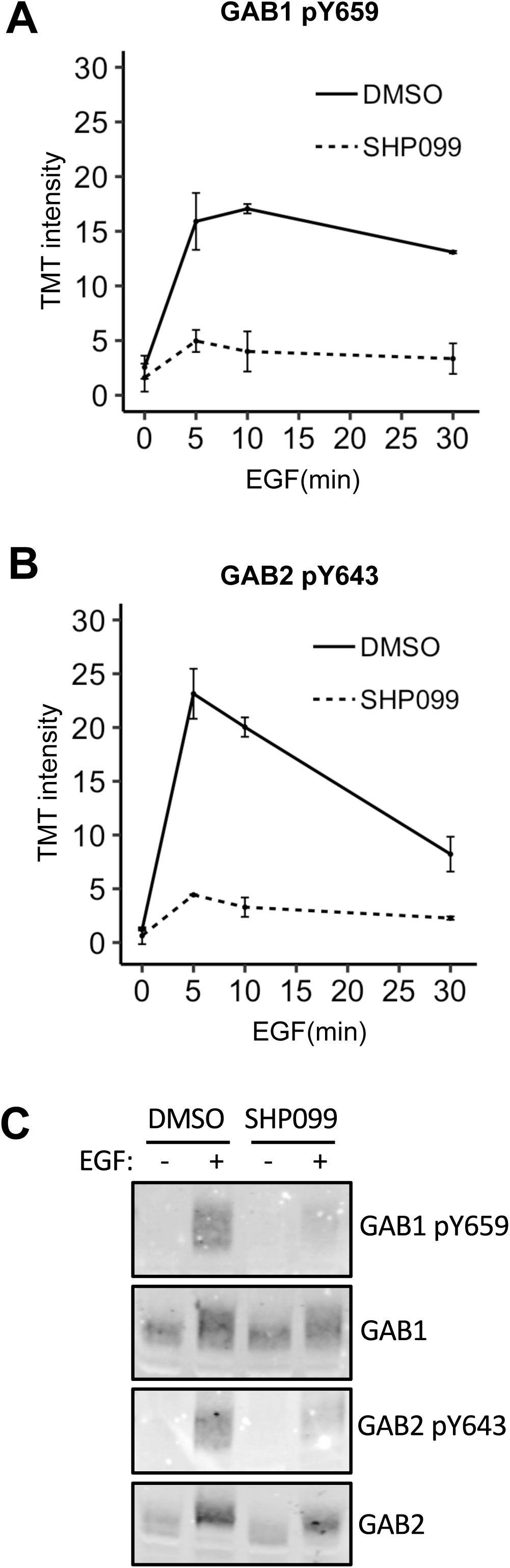
SHP2 inhibition results in reduced pY abundance at its interaction motifs. (A, B) TMT signal-to-noise intensities of GAB1 pY659 (A) and GAB2 pY643 (B) peptides showing dynamic changes in phosphorylation under DMSO- (solid line) and SHP099-treated (dotted line) conditions. (C) MDA-MB-468 cells pre-treated with DMSO carrier or SHP099 (10 µM) for 2 h were mock-treated or stimulated with EGF (10 nM) for 10 min. Total GAB1, total GAB2, GAB1 pY659 and GAB2 pY643 were analyzed by Western blot with both anti-protein and phosphospecific antibodies.

To determine whether pY sites of GAB1 and GAB2 exhibit the same pattern of response in other cellular contexts, we also treated other EGFR-driven cancer cell lines with SHP099. KYSE520, an EGFR-amplified esophageal cancer cell line, showed loss of GAB1 Y659 phosphorylation with SHP099 treatment under conditions of EGF stimulation (**Figure S3A**). H1975 cells, which carry an EGFR activating mutation, displayed high levels of constitutive GAB2-Y643 phosphorylation. SHP099 treatment eliminated phosphorylation of this site, and washout of the drug restores the mark within 5 minutes (**Figure S3B**). In addition, we tested whether deletion of the SHP2 gene recapitulated the effects of chemical inhibition on the phosphorylation of Y659 of GAB1. For these studies, we used a *PTPN11* knockout U2OS cell line prepared by CRISPR/Cas9 mediated genome editing (LaRochelle et al., 2018). When stimulated with EGF, SHP2-null U2OS cells showed a reduction in GAB1 pY659 levels when compared to parental cells, and reintroduction of SHP2 fully rescued the level of accumulated pY659 (**Figure S3C**). Stimulation of U2OS cells with PDGFββ, also showed an induction of the GAB1 pY659 mark that is lost upon treatment with SHP099 (**Figure S3D**). Similarly, stimulation of the T cell receptor on Jurkat cells with an anti-CD3 antibody induced GAB1 and GAB2 phosphorylation, and this induction was attenuated by SHP099 treatment (**Figure S3E**). These findings show that the effect of SHP2 on the abundance of GAB1 and GAB2 phosphotyrosine marks is broadly shared among a range of growth factor and antigen receptor signaling systems.

To determine how inhibition of SHP2 reduced phosphotyrosine abundance at these sites, we generated a set of site specific SHP2 mutants and asked whether they could rescue GAB1-pY659 phosphorylation in U2OS *PTPN11* null cells. We constructed point mutations that abolish autoinhibition (E76K), eliminate catalytic activity (C459E), or disable both autoinhibition and catalytic activity (E76K/C459E). We also created protein truncations that contain only the catalytic domain (PTP) or only the SH2 domains (SH2) (**Figure 5A**). When stimulated with EGF, C459E, E76K, and the E76K/C459E double mutant all restored GAB1-pY659 phosphorylation to a level similar to that of cells rescued with wild-type SHP2 (**Figure 5B**). When the SH2 or PTP constructs were introduced into U2OS cells lacking endogenous SHP2, the PTP fragment failed to rescue, whereas the SH2 fragment was as active as wild-type SHP2.

**Figure 5.**
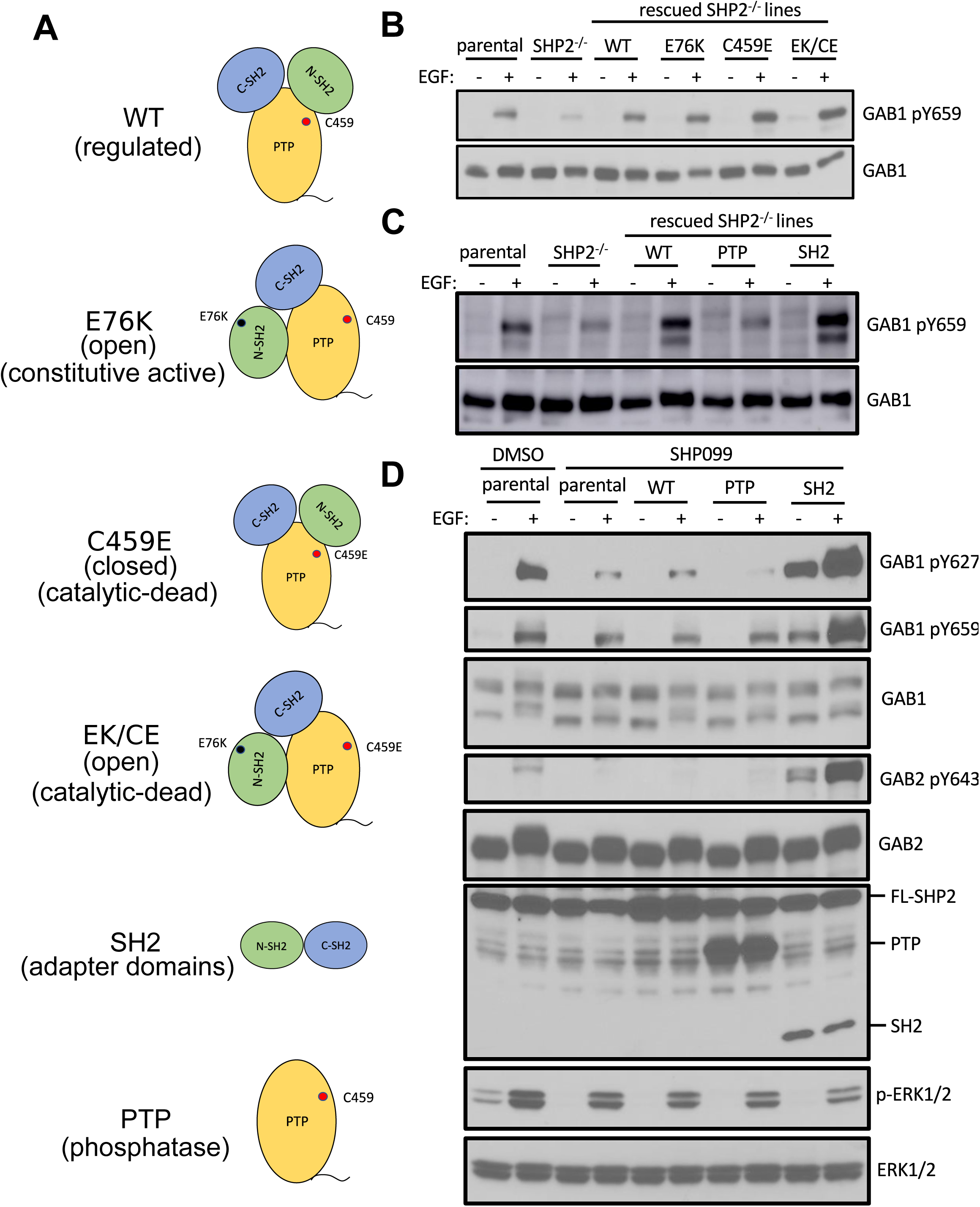
The SH2 domains of SHP2, but not its PTP domain, protect specific GAB1 and GAB2 pY marks. (A) Schematic representation of wild-type, point-mutants, and deletion constructs of SHP2 tested. (B, C) Parental-, SHP2 knockout- or SHP2 knockout U2OS cells stably expressing various SHP2 “rescue” constructs were cultured with or without EGF stimulation (10 nM) for 10 min. Cells were lysed and subjected to Western blotting using anti-pY659-GAB1 and anti-GAB1 antibodies. (D) MDA-MB-468 cells stably expressing wild-type, PTP or SH2 domains pre-treated with SHP099 were stimulated with EGF and immunoblotted after lysis using the specified antibodies.

Since MDA-MB-468 cells cannot survive without SHP2 (based on the gene essentiality data from avana_public_18Q2 library, Project Achilles (Meyers et al., 2017)), we stably expressed the SH2 or PTP fragments in parental MDA-MB-468 cells and chemically inhibited endogenous SHP2. These cell lines were stimulated with EGF and probed for phospho-GAB1 and phospho-GAB2. As expected, SHP099 treatment significantly reduced the abundance of pY659 on GAB1, as well as the abundance of pY643 on GAB2. GAB1 pY627, which has been reported to bind to the N-SH2 domain of SHP2 (Cunnick et al., 2001), also showed reduced phosphorylation (the corresponding mark on GAB2 (pY614) was not tested because commercial antibodies that detect this site are not available). Consistent with results from U2OS cells, the SH2 fragment, but not the PTP fragment was active in increasing pY abundance at these positions of GAB1 and GAB2 (**Figure 5D**). Together, these findings show that the catalytic activity of SHP2 is not required for the protection of the pY659 and pY643 marks on GAB1 and GAB2, respectively, and argue that the presence of tandem SH2 domains in SHP2 is needed to bind and protect phosphotyrosines on GAB1 and GAB2 from phosphatase-mediated dephosphorylation.

To assess whether this protection mechanism applies to other SHP2 interacting proteins that show reduced pY abundance in the presence of SHP099, we studied MPZL1, a cell surface receptor involved in extracellular matrix-induced signaling (Beigbeder, Chartier, & Bisson, 2017; Zhao, Guerrah, Tang, & Zhao, 2002). MPZL1 contains an immunoreceptor tyrosine-based inhibitory motif (ITIM) that, when doubly phosphorylated at Y241 and Y263, can bind the tandem SH2 domains of SHP2 (Zhao & Zhao, 2000). Our phosphoproteomic data show that basal phosphotyrosine abundance at these sites is greatly reduced by SHP099 treatment and that the dependence of pY modifications on SHP2 at these two sites occurs independent of EGF (**Figure S4A,B**). Using immunoblot assays, we found that the tandem SH2 domains alone are also sufficient to protect the phosphate groups on Y241 and Y263 of MPZL1 (**Figure S4C**).

Multiple studies have reported that GAB1, GAB2, and MPZL1 can be phosphorylated by Src family kinases (SFKs) (Chan et al., 2003; Kong, Mounier, Dumas, & Posner, 2003; Kusano, Thomas, & Fujiwara, 2008). Indeed, MDA-MB-468 cells treated with an SFK inhibitor (saracatinib/AZD0530) showed greatly reduced modification of GAB1-Y659 and GAB2-Y643 (**Figure S5A),** suggesting that SFK activity is required to phosphorylate the SHP2 binding sites on GAB1 and GAB2. SHP099, however, had no detectable effect on the abundance of phosphotyrosine at Y416 of the activation loop of SFKs (**Figure S5B)**, nor on the abundance of phosphotyrosine at Y527, which maintains SFKs in their autoinhibitory conformation (Roskoski, 2005). Together with the phosphosite protection results from the studies with various forms of SHP2 above, these data suggest that SHP2 acts to increase the half-life of phosphotyrosine modifications at sites on GAB1 and GAB2 that are phosphorylated by SFKs and then bound by SHP2.

### SHP2 is required for membrane localization of GAB1

To determine whether activation of SHP2 is required for recruitment of GAB1 to EGFR at the plasma membrane, we used a proximity ligation assay (PLA). MDA-MB-468 cells were mock-treated or stimulated with EGF in the presence of DMSO or SHP099, fixed in paraformaldehyde and probed for co-localization of GAB1 and EGFR. Under serum-starved (basal) conditions, the PLA signal was minimal and unaffected by SHP2 inhibition with SHP099. In contrast, within 2 minutes of EGF addition, cells treated with DMSO alone displayed a strong PLA signal indicative of GAB1 and EGFR co-localization, whereas SHP099 treatment reduced the number of binding events by 85% (p < 10^−4^; **Figure 6**). These results suggest that SHP2 is required to assemble the EGFR-GAB1 signaling hub crucial for MAPK and PI3K signaling.

**Figure 6.**
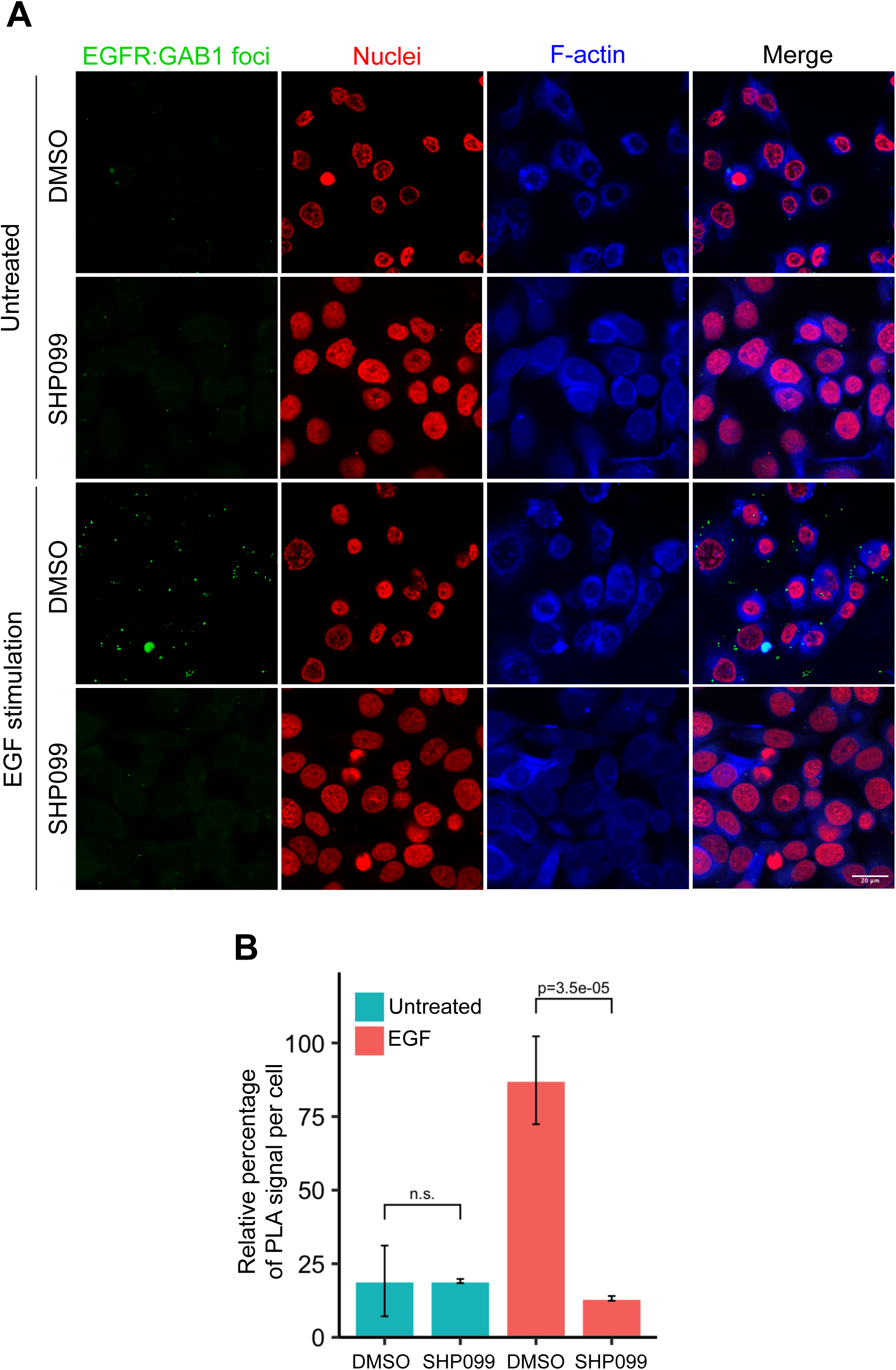
SHP2 regulates GAB1 association with activated EGFR. (A) Confocal images of EGFR-GAB1 complexes in MDA-MB-468 cells. Cells were pre-treated with DMSO carrier or SHP099 for 2 h, mock-treated or stimulated with 1 nM EGF for 2 min, and then analyzed using a proximity ligation assay (PLA). Cells were identified with propidium iodide (nuclei) and phalloidin (F-actin). PLA foci appear as green dots. Images are representative of three independent experiments. (B) Plot of relative PLA signal per cell (mean ± SD), quantifying the extent of EGFR-GAB1 complex formation under each treatment condition in (A).

## Discussion

Here, we performed quantitative phosphoproteomics to determine how the responses of cells to EGFR stimulation are modulated by inhibition of SHP2. We identified over 400 pY sites that exhibit increases or decreases in phosphorylation in response to SHP099 addition and washout. The dynamics of these changes in pY abundance reveal six distinct response signatures that provide a number of insights into how cancer cells depend on SHP2 in RTK signalling. Three of the six patterns identified by hierarchical clustering correspond to EGFR-dependent phosphotyrosine abundance changes that are also modulated by SHP2. These patterns include an early SHP2 substrate-like pattern (early responders that increase in abudnace upon SHP2 inhibition), a late SHP2 substrate-like pattern (late responders also increased upon SH2 inhibition), and a pattern in which phosphotyrosine levels are unexpectedly decreased when SHP2 is inhibited (protected sites).

Certain early responder sites lie on proteins that transmit information from activated RTKs to Ras/MAPK and PI3K signaling cascades. For example, the pY209 mark on the adaptor protein Grb2, which inhibits Sos binding and downstream Ras activation (Chardin et al., 1993; Li, Couvillon, Brasher, & Van Etten, 2001), rapidly accumulates upon SHP2 inhibition, suggesting that one important role of SHP2 in these cells is to remove an inhibitory mark and promote Grb2-dependent protein-protein interactions that drive immediate early signaling (**Figure 7**). Similarly, the adapter Crk also has two early responder pY sites at positions 108 and 136 that accumulate upon SHP2 inhibition; their removal by SHP2 may promote the ability of Crk to propagate the EGFR signal. Likewise, the PI3K regulatory subunits PI3KR1 and PI3KR2 accumulate pY marks in their inter-SH2 (iSH2) domains at analogous positions between residues 452-470 that may suppress effector function when SHP2 is inhibited, and PI3KR3 also has a pY residue in its iSH2 domain that also shows a substrate-like response at later timepoints. Not surprisingly, SHP2 inhibition also affects the pY abundance on numerous proteins involved in cell adhesion and migration, as well as on cytoskeletal components (Figure 3D).

**Figure 7.**
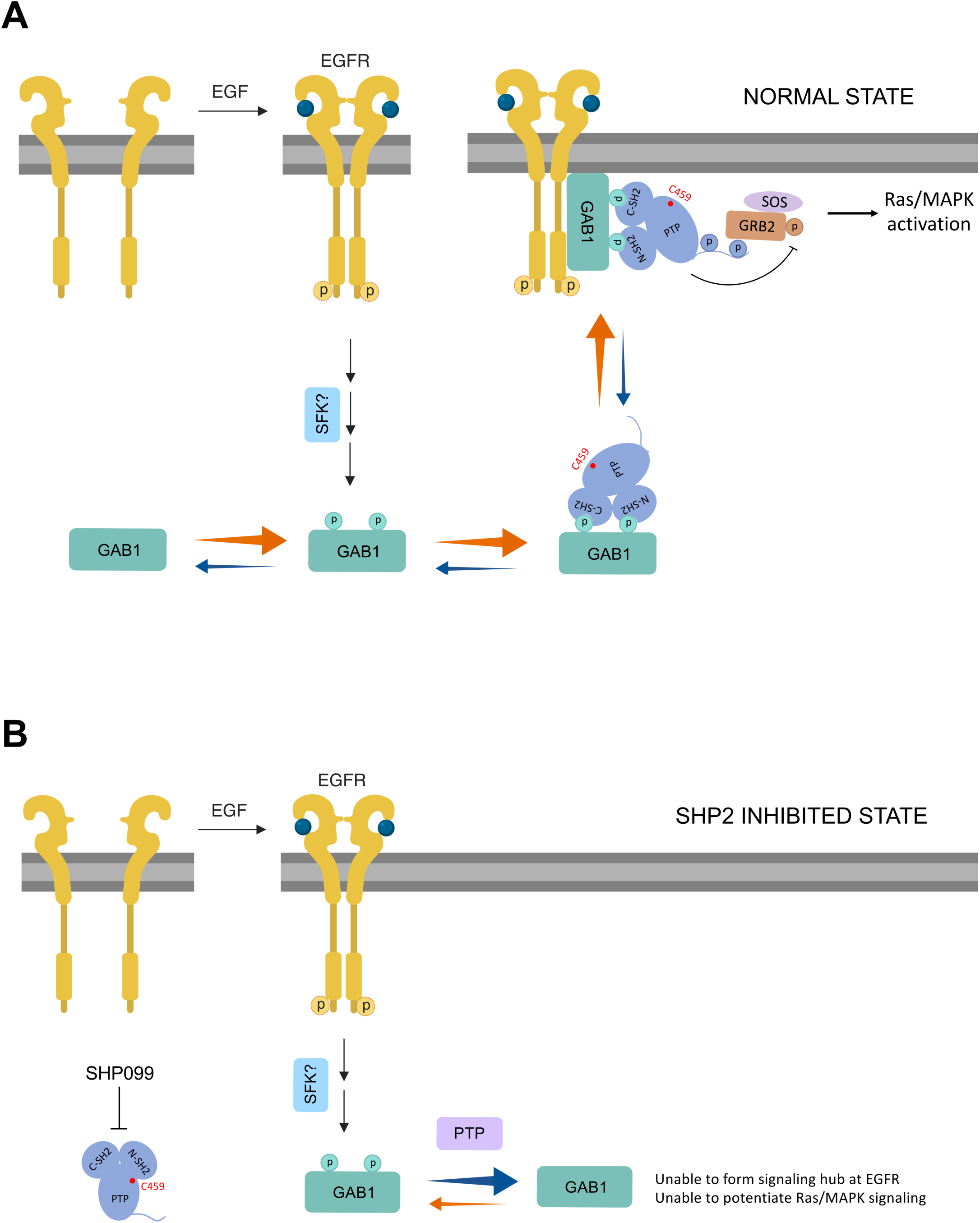
Model for cooperation of SHP2 with GAB1 in Ras/MAPK signal induction. (A) Upon growth factor stimulation, wild-type SHP2 stabilizes an EGFR-GAB1-GRB2-SOS signaling hub to potentiate Ras/MAPK activation. (B) In the presence of SHP099, SHP2 cannot form complexes with phosphorylated GAB1, preventing membrane recruitment and formation of a stable EGFR signaling hub.

Our data also reveal many proteins with SHP2-dependent changes in pY abundance that function in cellular processes previously unlinked to SHP2. These proteins include effectors of distal signaling responses such as phospholipase C enzymes (PLCγ1 and PLCγ2). Also noteworthy are the late-response, substrate-like patterns that predominate among proteins implicated in mRNA processing and endocytosis factors (hnRNPA2B1, hnRNPA3, EPN1/2, RAB10, FNBP1L, ITSN2, STX4) (**Figure 3D**). Dissecting the role of SHP2 in these events should be a fertile area of future investigation.

Strikingly, the most frequent change in pY pattern following SHP2 inhibition is not substrate-like, but rather one in which pY levels fall. Others have also recently reported that binding of SHP2 can prevent dephosphorylation of certain pY sites in the response to stimulation of other RTKs (Batth et al., 2018). This pattern of response is seen at pY residues on proteins recruited to the EGFR signaling hub, such as GAB1, GAB2, as well as on EGFR and SHP2 itself. To better understand this mode of pY regulation, we studied SHP2-dependent protection of pY sites on GAB1 and GAB2, adapters that induce SHP2 binding and activation upon tyrosine phosphorylation (Arnaud et al., 2004; Crouin, Arnaud, Gesbert, Camonis, & Bertoglio, 2001; Cunnick et al., 2001). Binding and mutagenesis studies clearly demonstrate that the tandem SH2 domains of SHP2 shield pY binding sites on GAB1 and GAB2 from dephosphorylation by SHP2 and presumably also other phosphatases. Also intriguing is the SHP2-mediated protection of nearby pairs of pY residues on several proteins, including MPZL1-pY241/pY263, a fibronectin-activated adhesion protein that can assemble into complexes containing SHP2 and Grb2 in a pY-dependent manner (Beigbeder et al., 2017). The protection of phosphotyrosine sites by SHP2 highlights the scaffolding role of SHP2 in stimulating phosphotyrosine engagement by SH2 domains at sites of RTK activation. SHP2 scaffolding activity is particularly relevant to Noonan and LEOPARD syndromes, both of which result from missense SHP2 mutations. Though distinct, the two syndromes have overlapping clinical features. Mutations found in Noonan syndrome destabilize the autoinhibited form of the enzyme without substantially affecting the intrinsic catalytic activity of the phosphatase domain, leading to increased basal phosphatase activity, elevated scaffolding activity, and downstream induction of ERK (Araki et al., 2004; Niihori et al., 2005). LEOPARD syndrome mutations also destabilize the autoinhibited conformation of the enzyme, enhancing scaffolding activity, but in addition cripple the catalytic activity of the phosphatase domain. The increased preference for the open conformation in both Noonan and LEOPARD syndrome mutations promotes the protective “arm” of SHP2 signaling, whereas only Noonan syndrome mutations retain the catalytic activity needed to drive the substrate-dependent features of signaling. The work reported here shows how central the scaffolding function is at the EGFR signaling hub and in the engagement of pY sites on GAB1 and GAB2 at the site of activation, with the phosphatase-dependent events, through adapter proteins like Grb2 and key signaling nodes like PI3K, required for full propagation of a growth factor induced signal to downstream effectors.

## Materials and Methods

### Plasmids, compounds and ligands

Constructs used to rescue SHP2 knockout U2OS cells were obtained by cloning the SHP2 cDNA from pCMV-SHP2-WT (Addgene plasmid # 8381) into the migR1-IRES-GFP vector (Addgene plasmid # 27490). Point mutants (E76K, C459E, and DM) were generated from the parental SHP2-migR1 construct using site-directed mutagenesis (QuikChange II, Agilent). SH2-only and phosphatase-only mutants were generated by cloning the PTP (220-525 aa) and N-SH2-C-SH2 (1-219 aa) fragments, respectively, into the migR1-IRES-GFP vector. SHP099 was obtained commercially from DC chemicals (Catalog No. DC9737). AZD0530 (Saracatinib) was a gift from Dr. Nathanael Gray (Dana-Farber Cancer Institute). The CD3 monoclonal antibody (UCHT1) was purchased from Thermo Fisher Scientific (Catalog No. 16-0038-85). Recombinant human EGF and PDGF-BB were purchased from Gibco (Catalog No. PHG0311) and Peprotech (Catalog No. 10771-922), respectively.

### Antibodies

Antibodies used in this study were obtained commercially from the following sources: Phospho-Tyr-1000 (CST, #8954), Phospho-Thr202/Tyr204-Erk1/2 (CST, #9101), Erk1/2 (CST, #9102), GAB1-pY659 (CST, #12745), GAB1-pY627 (CST, #3231), GAB2-pY643 (Thermo Fisher Scientific, #PA5-37778), GAB1 (CST, #3232), GAB2 (CST, #3239), pY263-MPZL1 (CST, #5543), pY241-MPZL1 (CST, #8131), MPZL1 (CST, #4157), pY1005-RhoGAP (Thermo Fisher Scientific, PA5-36713), RhoGAP (CST, #2562) and SHP2 (CST, #3397).

### Cell culture and generation of stable cell lines

All the parental cell lines used in this study (MDA-MB-468, KYSE520, U2OS, Jurkat and NCI-H1975) were purchased from ATCC. The SHP2 knockout U2OS cell line was created previously (LaRochelle et al., 2018). To generate rescued SHP2-null U2OS lines, knockout cells were infected with retrovirus harboring the wild-type or mutant SHP2 cDNAs and FACS sorted for GFP positive cells. MDA-MB-468 parental cells stably expressing full-length GAB1-GFP, GAB2-GFP, SHP2, PTP and N-SH2-C-SH2 cDNAs were also generated similarly. U2OS cell lines were grown in McCoy’s 5A media with 10% FBS. MDA-MB-468 cells were cultured in Leibovitz’s L-15 media with 10% FBS without CO2. All other cell lines (KYSE520, Jurkat and NCI-H1975) were maintained in RPMI-1640 supplemented with 10% FBS at 5% CO2.

### Ligand stimulation

For ligand stimulation experiments analyzed by Western blotting, cells were seeded at 70% confluence in 10 cm petri dishes (Nunc, Thermo Fisher Scientific), serum starved for 24h, and treated with DMSO carrier or SHP099 (10 uM) for 2h before stimulation with ligands (EGF (10 nM), PDGF-BB (5 nM) or anti-CD3 (10 ug/ml)) for the indicated time periods. Cells were then quickly washed with ice-cold PBS twice, lysed in Ripa buffer (25 mM Tris-HCl (pH 7.6), 150 mM NaCl, 1% Nonidet P-40, 1% Sodium deoxycholate, 0.1% Sodium dodecyl sulfate, 2 mM EDTA) with protease and phosphatase inhibitors (cOmplete Mini and PhosSTOP, Roche) and analyzed by Western blotting.

### Sample preparation for phosphoproteomics

MDA-MB-468 cells were seeded in 15 cm cell culture dishes (Corning) in full growth media (Leibovitz’s L-15 with 10% FBS and 1% penicillin/streptomycin). At 80% confluence, cells were washed twice with warm HBSS buffer and incubated in serum-free media for 24h. On the following day, cells were pre-treated with DMSO or SHP099 (10 µM) for 2h prior to ligand stimulation. As outlined in Figure 1A, DMSO and SHP099 groups were left untreated (0 min) or stimulated with EGF (10 nM) for 5, 10 or 30 min. The SHP099 washout group was stimulated with EGF (10 nM) for 10 min, washed three times with warm HBSS and stimulated with EGF again for 5, 10 or 30 min. To terminate stimulation, cells were immediately washed with ice-cold PBS, harvested, centrifuged, flash frozen in liquid N2 and stored at −80°C until all 11 samples were prepared. Cell pellets were lysed in lysis buffer (2% SDS, 150 mM NaCl, 50mM Tris (pH 8.5-8.8)) containing protease and phosphatase inhibitors (cOmplete Mini, PhosSTOP, Roche; 2mM Sodium Orthovanadate, NEB), sonicated to shear chromatin and centrifuged to remove cellular debris. A bicinchoninic acid (Pierce) assay was performed according to manufacturer’s instructions and lysates were normalized to a protein concentration of 1 mg/ml, reduced with DTT (5 mM), alkylated with iodoacetamide (14 mM) and quenched with further addition of DTT (5 mM). Total protein (1 mg) was precipitated with methanol-chloroform, reconstituted in 8 M urea at pH 8.5, sonicated, diluted to 4 M urea and digested with LysC overnight and Trypsin for 6 h at an enzyme-to-protein ratio of 1:100. The percentage of missed cleavages was monitored using aliquots from a few representative samples (total 3 µg) that were desalted and analyzed by mass spectrometry. Digests were acidified with 2% formic acid and desalted using C18 Sep-Pak cartridges (WAT054960, Waters). Samples were dried by vacuum centrifugation, resuspended in 80% acetonitrile/0.15% trifluoroacetic acid (TFA) and normalized to 1 mg/ml for enrichment of phosphopeptides.

### Phosphopeptide enrichment

Phosphopeptides were enriched using Immobilized Metal Affinity Chromatography (IMAC) with NTA Magnetic Agarose beads (Qiagen, Cat # 36113) stripped of Ni^2+^ and reloaded with Fe(III). 500 µl of beads were incubated in 1 ml EDTA (40 mM) for 30 min at room temperature to remove bound Ni^2+^, followed by three washes with 1 ml of HPLC-grade water. The beads were then charged with iron using FeCl_3_ (100 mM) for 30 min at room temperature. Excess iron was removed by washing the beads three times with water followed by acidification with 80% acetonitrile/0.15% TFA. Peptides (1 mg per sample) were incubated with the IMAC beads for 30 min at room temperature. Phosphopeptide bound beads were washed with 1 ml of 80% acetonitrile/0.15% TFA to remove non-specifically bound peptides. Phosphopeptides were eluted with 300 µl of 50% acetonitrile/0.7% NH_4_OH, acidified with 4% formic acid, immediately vacuum centrifuged to dryness and subjected to desalting using SOLA HRP 10mg Sep-Pak cartridges (Thermo-Fisher).

### Tandem Mass Tag (TMT) labeling

Prior to isobaric labeling, phosphopeptides were reconstituted in 200 mM EPPS (pH 8.3) and anhydrous acetonitrile to 30% (v/v). Each sample was labeled with 5 µl of a TMT 11-plex reagent (20 µg/µL; A37725, Thermo Scientific) for 90 min at room temperature. The reactions were quenched with 5 µL of 5% hydroxylamine for 15 min, combined, acidified with TFA to 1.0% (v/v), desalted using 10 mg SOLA Sep-Pak cartridges (Thermo Fisher) and lyophilized for 2 days to remove residual TFA.

### Phosphotyrosine Immunoaffinity Purification (pY IAP)

The phosphotyrosine antibody (p-Tyr-1000, Cell Signaling Technology) was coupled with agarose beads one day prior to the IAP as follows. A Protein A agarose bead slurry (Sigma-Roche) (60ul) was washed four times with 1ml of cold PBS. The p-Tyr-1000 antibody (Cell Signaling Technology) (30ul) was gently mixed with 1.5ml PBS and added onto the washed agarose beads and incubated on a rotator overnight at 4°C. The beads were washed four times in 1ml of cold PBS to remove excess uncoupled antibody. Lyophilized peptides were reconstituted with 500ul of IAP buffer (50mM MOPS/NaOH pH 7.2, 10mM Na_2_HPO_4_, 50mM NaCl) and incubated with antibody conjugated beads for 2h at 4°C with gentle rotation. The nonspecific binding phosphopeptide (phospho-serine and -threonine peptides) flow through was collected after centrifugation and stored at −80°C. The phosphotyrosine bound peptides were washed once with 1ml of IAP buffer, transferred into a 0.2uM filter spin column followed by two 400 ul washes with cold HPLC-grade water. Phosphotyrosine peptides were eluted twice with 75ul of 100 mM formic acid. The pY eluate was desalted using a C18 StageTip, dried by vacuum centrifugation and reconstituted in 3% acetonitrile/0.5% formic acid for MS analysis.

### Mass Spectrometry

Phosphoproteomic mass spectrometric data were acquired on an Orbitrap Fusion Lumos mass spectrometer (Thermo Fisher Scientific) equipped with a Proxeon EASY-nLC 1000 liquid chromatography system (Thermo Fisher Scientific). Phosphopeptides were separated on a 75 µm inner diameter microcapillary column packed with ∼35 cm Sepax GP-C18 resin (1.8 µm, 150 A, Thermo Fisher Scientific). Phosphopeptides (∼2 µg) were separated using a 2h gradient of acetonitrile in 0.125% formic acid.

The phosphoproteome MS analysis scan sequence began with the collection of an FTMS1 spectrum (120,000 resolution with mass range 400-1400 Th). The top 10 most intense ions were selected for MS/MS and fragmented via collision-induced dissociation (CID, CE = 35%) with a maximum injection time of 200 ms and an isolation window of 0.5 Da. FTMS3 precursors were fragmented by high energy collision-induced dissociation (HCD, CE = 55%) and analyzed at 50,000 resolution (200 Th) with a maximum ion injection time of 300 ms and an isolation window of 1.2. The MultiNotch MS3-based TMT method was applied as described in McAlister et al (McAlister et al., 2014).

### Data processing

Mass spectra were processed using a SEQUEST-based software pipeline. A modified version of ReAdW.exe was used to convert spectra (.raw) to mzXML. All spectra were searched against a database containing the human proteome downloaded from Uniprot (February 4th 2014) and common contaminating protein sequences. Database searches were performed using a peptide mass tolerance of 50 ppm and a fragment ion tolerance of 0.9 Da. Tandem mass tags (TMT) on lysine residues and peptide N termini (+229.163 Da) and carbamidomethylation of cysteine residues (+57.021 Da) were fixed modifications, while oxidation of methionine residues (+15.995 Da) and phosphorylation of serine, threonine, and tyrosine residues were set as a variable modification (+79.966 Da).

Peptide-spectrum matches (PSMs) were filtered by linear discriminant analysis to a false discovery rate (FDR) of 2% at the peptide level based on matches to reversed sequences. Linear discriminant analysis considered the following parameters: XCorr, ΔCn, missed cleavages, adjusted PPM, peptide length, fraction of ions matched, charge state, and precursor mass accuracy. Filtered peptides were collapsed further to a final protein-level FDR of 2%. Phosphopeptides were quantified from MS3 scans after filtering with a total TMT reporter signal-to-noise ratio > 200 and isolation specificity at 0.5. Localization of phosphorylation sites was determined using AScore (Huttlin, E.L. Cell 2010), and sites with an AScore > 13 were selected for further analysis.

### Dependence classification of phosphorylation site time courses

Each phosphorylation time course was classified according to its response to 1) EGF stimulation and 2) SHP2 inhibition. This analysis was performed using the union of the two biological replicates, averaging measurements for sites that were found in the intersection of replicates. EGF classes were defined based on phosphorylation dynamics in control cells only. Significant increases and decreases were defined as changes of at least 1.5-fold relative to time 0 (unstimulated cells). The classes comprised the following: 1) fast increase (increased phosphorylation within 5 minutes of EGF stimulation), 2) medium increase (increased phosphrylation within 10 minutes), 3) slow increase (increased phosphorylation within 30 minutes), 4) neutral (no change at any timepoint), and 5) decrease (decreased phosphorylation by any point in the time course).

#### EGF classes

| **Name** | **Earliest timepoint where change is detected** |
| --- | --- |
| Fast increase | 5 min |
| Medium increase | 10 min |
| Slow increase | 30 min |
| Neutral | n/a |
| Decrease | Any |

Classes of SHP2 responses were defined based on comparison between control cells and cells treated with SHP099 for two hours prior to EGF stimulation. The classes comprised the following: 1) pre-stimulation negative (increased phosphorlyation in SHP099-treated cells compared to control cells, prior to EGF addition), 2) post-stimulation negative (increased phosphorylation in SHP099-treated cells at any point after EGF addition), 3) neutral (no difference between treatment and control at any timepoint), 4) pre-stimulation positive (decreased phosphorylation in SHP099-treated cells compared to control cells, prior to EGF addition), and 5) post-stimulation positive (decreased phosphorylation in SHP099-treated cells at any point after EGF stimulation).

#### SHP2 classes

| **Name** | **Change with SHP099 treatment** | **Timing of change** |
| --- | --- | --- |
| Pre-stimulation negative | Increase | Before EGF addition |
| Post-stimulation negative | Increase | Any timepoint after EGF addition, but not before |
| Neutral | None | n/a |
| Pre-stimulation positive | Decrease | Before EGF addition |
| Post-stimulation positive | Decrease | Any timepoint after EGF addition, but not before |

### Bioinformatic analysis

Volcano plots were generated for all sites that were detected in either of the biological replicates. For each site at each timepoint (5 min, 10 min and 30 min), the mean of [SHP099/DMSO] and [SHP099/WO] fold changes was plotted against –log10(p-value). Significantly altered phosphorylation sites were visualized based on their membership in the Reactome databse (http://software.broadinstitute.org/gsea/msigdb/annotate.jsp). Gene set analysis was performed using publicly available GSEA tools (Mootha et al., 2003; Subramanian et al., 2005), which were used to compute the overlap between detected proteins and the REACTOME pathway database. Hierarchical clustering was performed using the clustergram function of MATLAB with a Euclidean distance metric.

### Proximity Ligation Assay

MDA-MB-468 cells were grown to 70% confluence on 8-chambered glass slides (Nunc Lab-Tek II, Thermo Scientific), starved in serum-free media for 24 h,and were treated with DMSO carrier or SHP099 (10 uM) for 2 h prior to stimulation with 1 nM EGF for 2 min. Cells were fixed with ice-cold 4% (v/v) paraformaldehyde for 15 minutes and permeabilized with 0.125% Tween-20 in PBS for 5 minutes prior to incubation in 1X blocking solution (1X PBS, 10% BSA, 10% donkey serum, 10% goat serum) for 1 h at room temperature. Cells were then incubated with mouse anti-EGFR (1:1000, clone 13G8, Millipore) and rabbit anti-GAB1 (1:100, catalog # SAB4501060, Sigma) overnight at 4°C in a humidified chamber. The proximity ligation assay was conducted using a DuoLink In Situ Mouse/Rabbit PLA kit (Sigma-Aldrich) according to the manufacturer’s instructions. All images were collected on a confocal laser scanning microscope (Zeiss LSM 880 with Airyscan) equipped with a 63x oil immersion objective lens (Plan Apo, NA=1.4), and analyzed using Cell Profiler software.

## Acknowledgments

This work was supported by NIH grants R35 CA220340 (SCB), and U54-HL127365 (PKS). We thank Jon Aster, Kimberly Stegmaier, Huaixiang Hao, Aram Ghalali, Abhinav Dubey and members of the Blacklow laboratory for helpful discussions.

---

**Figure S1.**
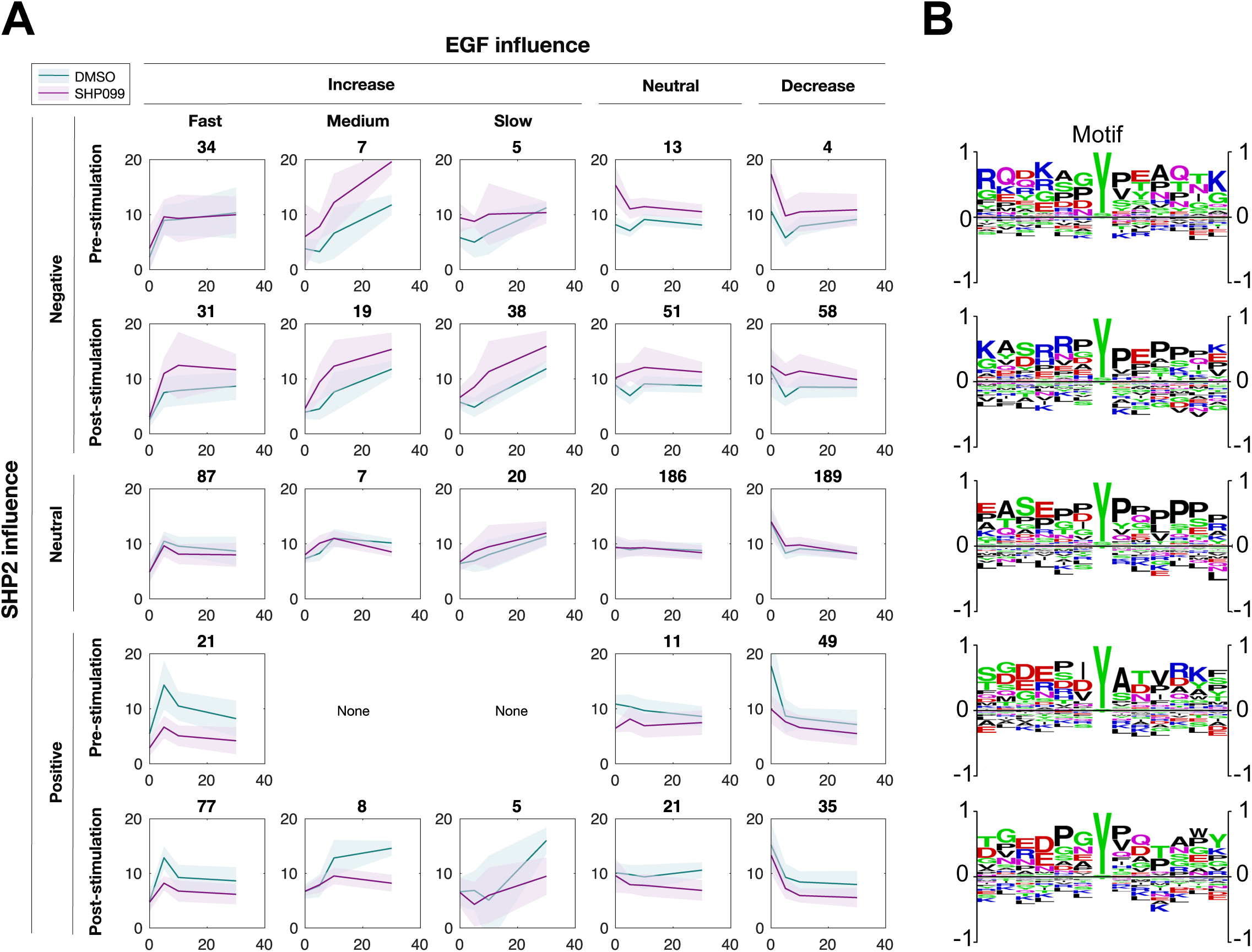
Global analysis of the SHP2 regulated phosphotyrosine-proteome. (A) Classification of phosphorylation time courses based on EGFR and SHP2 dependence. Numbers above each plot indicate the number of phosphopeptides within each class combination. (B) Sequence logos showing amino acid preferences at positions flanking the phosphorylated tyrosine residue for each cluster.

**Figure S2.**
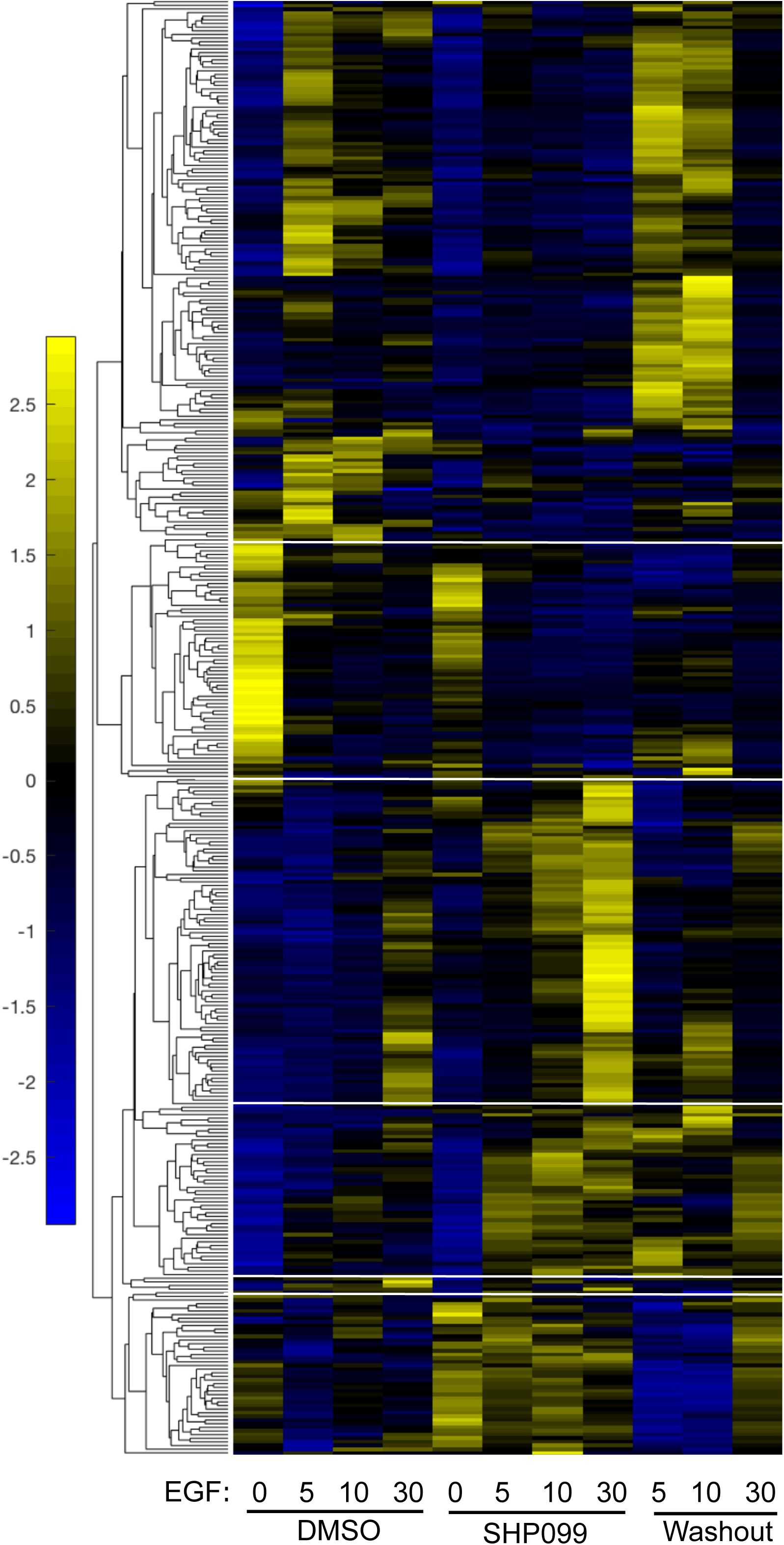
Kinetic pattern of pY response to EGF stimulation under DMSO, SHP099, and washout conditions. The heatmap shows six different response patterns, based on hierarchical clustering.

**Figure S3.**
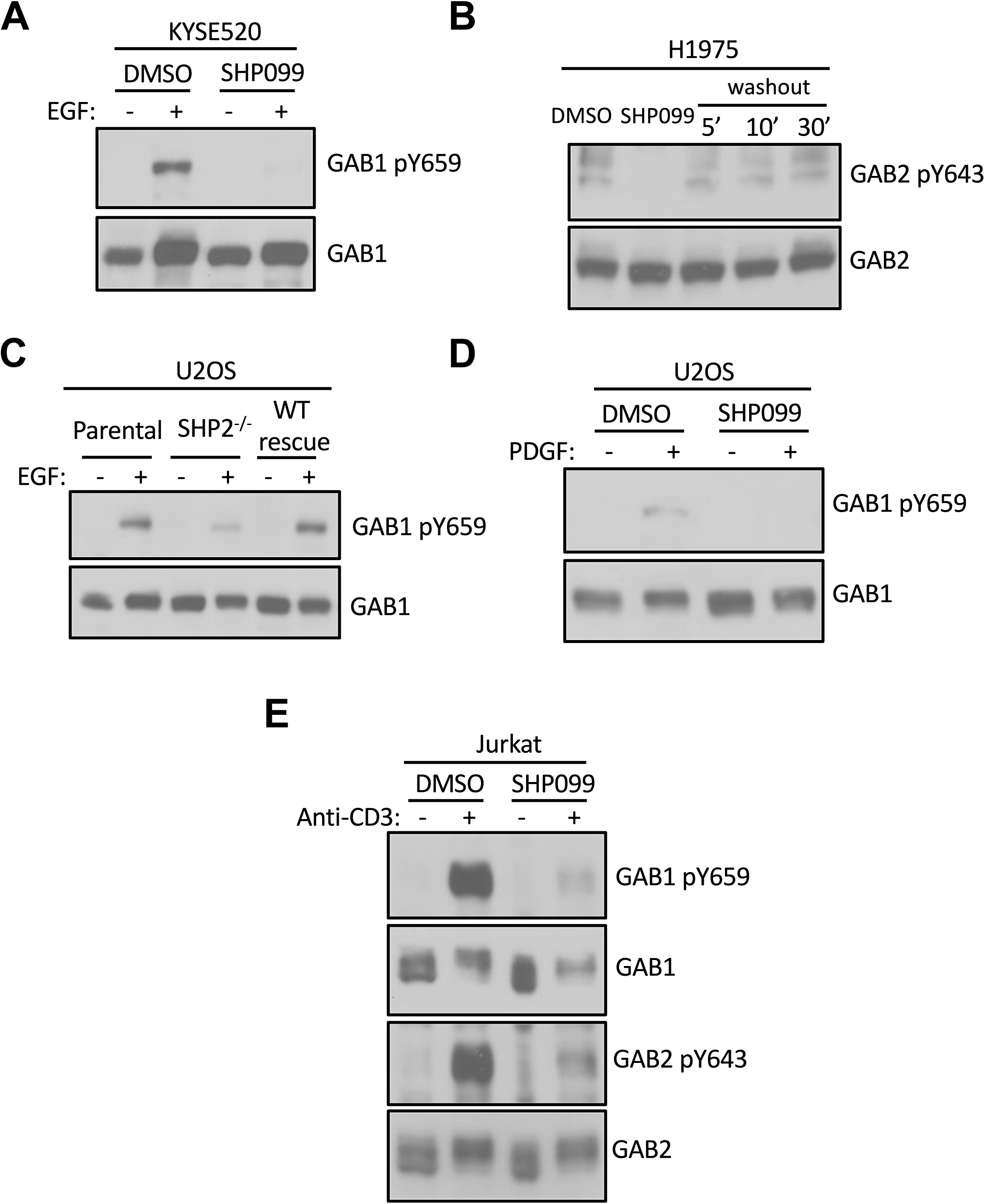
SHP2 regulates GAB1 and GAB2 phosphorylation in multiple cellular contexts. (A) KYSE520 cells pre-treated with DMSO carrier or SHP099 (10 uM) for 2 h were left uninduced or induced with EGF (10 nM) for 10 min. Whole cell lysates were then subjected to immunoblotting with GAB1-pY659 and GAB1 antibodies. (B) H1975 cells were treated with SHP099 (10uM) for 2h, washed three times with warm HBSS buffer and cultured in full growth medium (RPMI-1640 supplemented with 10% FBS) for the indicated time periods. Lysates were then blotted with pY643-GAB2 and GAB2 antibodies. (C) Whole cell extracts from parental-, SHP2 knockout- and SHP2 knockout stably expressing wild-type SHP2 U2OS cell lines were blotted with pY659-GAB1 and GAB1 antibodies. (D) DMSO- and SHP099-treated U2OS cells were stimulated with PDGF (5 nM) and probed for pY659-GAB1 and GAB1. (E) DMSO- and SHP099-treated Jurkat cells were stimulated with anti-CD3 antibody (10 ug/ml) and subjected to immunoblotting with pY659-GAB1, pY643-GAB2, GAB1 and GAB2 antibodies.

**Figure S4.**
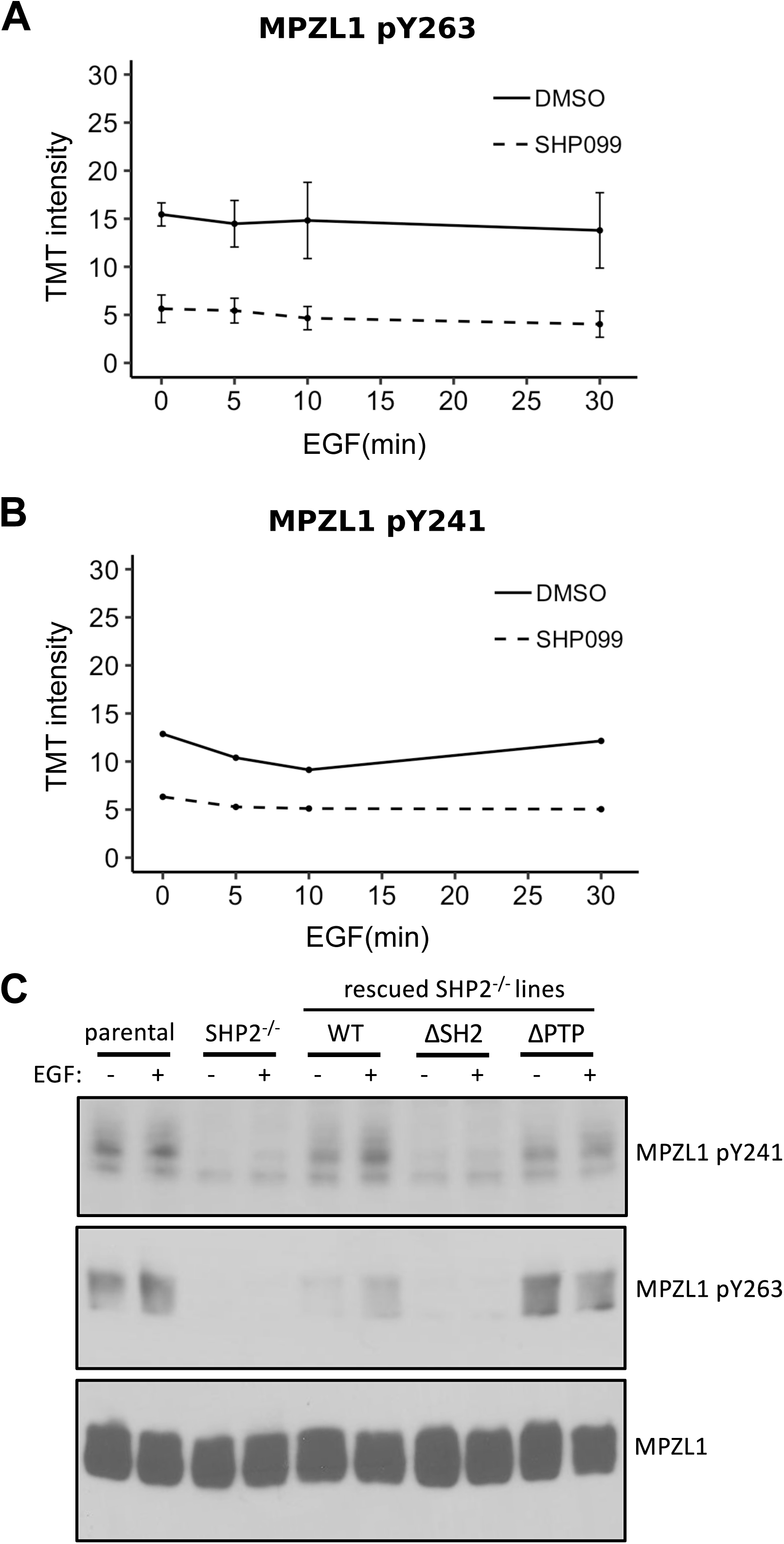
SHP2 binds and protects MPZL1 phosphorylation. (A and B) TMT signal-to-noise intensities of MPZL1 peptides showing phosphorylation changes between DMSO- (solid line) and SHP099-treated (dotted line) groups. (C) Unstimulated or EGF-stimulated parental-, SHP2 knockout- and SHP2 knockout stably expressing wild-type, PTP and SH2 U2OS cell lines were blotted for pY263-MPZL1, pY241-MPZL1 and MPZL1.

**Figure S5.**
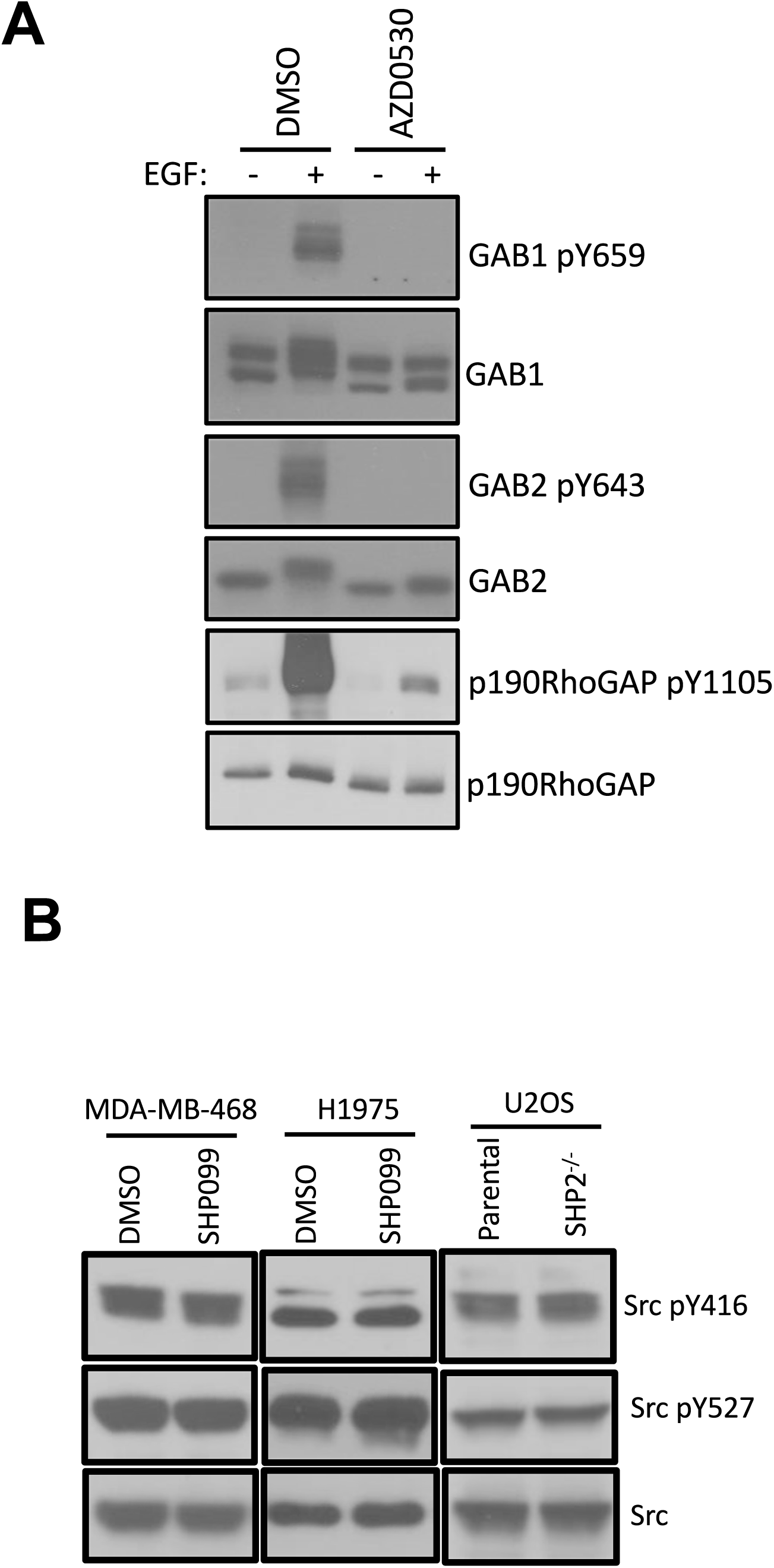
SHP2 does not affect the deposition of phosphate marks on GAB1/2 by SFKs. (A) MDA-MB-468 cells pre-treated with DMSO carrier or AZD0530 (10 uM) were left unstimulated or stimulated with EGF (10 nM) for 10 min and subjected to western analysis with pY659-GAB1 and pY643-GAB2 antibodies. pY1105-RhoGAP, a known substrate of Src, acts as a positive control for Src inhibition. Total protein levels were detected using GAB1, GAB2, RhoGAP antibodies. (B) MDA-MB-468 and H1975 cells were treated with DMSO and SHP099 (10uM) for 2h and subjected to immunoblotting with Src-pY416, Src-pY527 and Src antibodies. Parental and SHP2 deficient U2OS cells were probed for Src-pY416, Src-pY527 and total Src levels.

**Figure S6.**
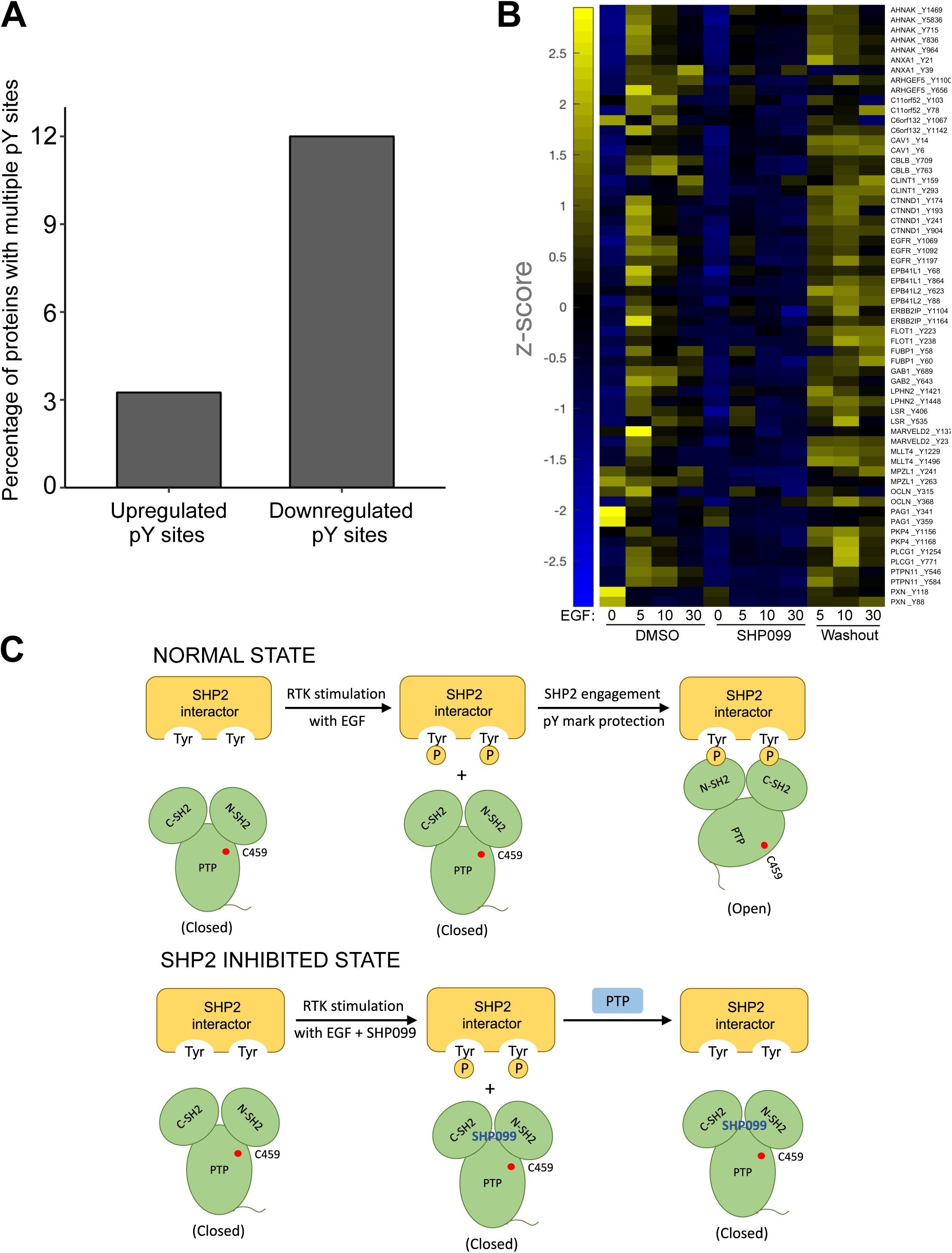
SHP2-protected pY sites are enriched for multisite protein phosphorylation. (A) Quantification showing the relative percentages of proteins displaying multisite phosphorylation in SHP099-responsive pY sites. (B) Heatmap visualization of multisite protein phosphorylation dynamics in the class of SHP2-protected pY sites. (C) Schematic model showing pY mark protection by the stable engagement with the tandem SH2 domains of SHP2. In the presence of SHP099, pY marks on putative SHP2 interactors are vulnerable to dephosphorylation.

## References

Agazie, Y. M., & Hayman, M. J. (2003). Molecular mechanism for a role of SHP2 in epidermal growth factor receptor signaling. Mol Cell Biol, 23(21), 7875–7886.

Araki, T., Mohi, M. G., Ismat, F. A., Bronson, R. T., Williams, I. R., Kutok, J. L., … Neel, B. G. (2004). Mouse model of Noonan syndrome reveals cell type-and gene dosage-dependent effects of Ptpn11 mutation. Nat Med, 10(8), 849–857. doi:10.1038/nm1084

Arnaud, M., Mzali, R., Gesbert, F., Crouin, C., Guenzi, C., Vermot-Desroches, C., … Bertoglio, J. (2004). Interaction of the tyrosine phosphatase SHP-2 with Gab2 regulates Rho-dependent activation of the c-fos serum response element by interleukin-2. Biochem J, 382(Pt 2), 545–556. doi:10.1042/BJ20040103

Batth, T. S., Papetti, M., Pfeiffer, A., Tollenaere, M. A. X., Francavilla, C., & Olsen, J. V. (2018). Large-Scale Phosphoproteomics Reveals Shp-2 Phosphatase-Dependent Regulators of Pdgf Receptor Signaling. Cell Rep, 22(10), 2784–2796. doi:10.1016/j.celrep.2018.02.038

Beigbeder, A., Chartier, F. J. M., & Bisson, N. (2017). MPZL1 forms a signalling complex with GRB2 adaptor and PTPN11 phosphatase in HER2-positive breast cancer cells. Sci Rep, 7(1), 11514. doi:10.1038/s41598-017-11876-9

Bennett, A. M., Tang, T. L., Sugimoto, S., Walsh, C. T., & Neel, B. G. (1994). Protein-tyrosine-phosphatase SHPTP2 couples platelet-derived growth factor receptor beta to Ras. Proc Natl Acad Sci U S A, 91(15), 7335–7339.

Bentires-Alj, M., Paez, J. G., David, F. S., Keilhack, H., Halmos, B., Naoki, K., … Neel, B. G. (2004). Activating mutations of the noonan syndrome-associated SHP2/PTPN11 gene in human solid tumors and adult acute myelogenous leukemia. Cancer Res, 64(24), 8816–8820. doi:10.1158/0008-5472.CAN-04-1923

Bregeon, J., Loirand, G., Pacaud, P., & Rolli-Derkinderen, M. (2009). Angiotensin II induces RhoA activation through SHP2-dependent dephosphorylation of the RhoGAP p190A in vascular smooth muscle cells. Am J Physiol Cell Physiol, 297(5), C1062–1070. doi:10.1152/ajpcell.00174.2009

Bunda, S., Burrell, K., Heir, P., Zeng, L., Alamsahebpour, A., Kano, Y., … Ohh, M. (2015). Inhibition of SHP2-mediated dephosphorylation of Ras suppresses oncogenesis. Nat Commun, 6, 8859. doi:10.1038/ncomms9859

Chan, P. C., Chen, Y. L., Cheng, C. H., Yu, K. C., Cary, L. A., Shu, K. H., … Chen, H. C. (2003). Src phosphorylates Grb2-associated binder 1 upon hepatocyte growth factor stimulation. J Biol Chem, 278(45), 44075–44082. doi:10.1074/jbc.M305745200

Chardin, P., Camonis, J. H., Gale, N. W., van Aelst, L., Schlessinger, J., Wigler, M. H., & Bar-Sagi, D. (1993). Human Sos1: a guanine nucleotide exchange factor for Ras that binds to GRB2. Science, 260(5112), 1338–1343.

Chen, Y. N., LaMarche, M. J., Chan, H. M., Fekkes, P., Garcia-Fortanet, J., Acker, M. G., … Fortin, P. D. (2016). Allosteric inhibition of SHP2 phosphatase inhibits cancers driven by receptor tyrosine kinases. Nature, 535(7610), 148–152. doi:10.1038/nature18621

Crouin, C., Arnaud, M., Gesbert, F., Camonis, J., & Bertoglio, J. (2001). A yeast two-hybrid study of human p97/Gab2 interactions with its SH2 domain-containing binding partners. FEBS Lett, 495(3), 148–153.

Cunnick, J. M., Mei, L., Doupnik, C. A., & Wu, J. (2001). Phosphotyrosines 627 and 659 of Gab1 constitute a bisphosphoryl tyrosine-based activation motif (BTAM) conferring binding and activation of SHP2. J Biol Chem, 276(26), 24380–24387. doi:10.1074/jbc.M010275200

Easton, J. B., Royer, A. R., & Middlemas, D. S. (2006). The protein tyrosine phosphatase, Shp2, is required for the complete activation of the RAS/MAPK pathway by brain-derived neurotrophic factor. J Neurochem, 97(3), 834–845. doi:10.1111/j.1471-4159.2006.03789.x

Garcia Fortanet, J., Chen, C. H., Chen, Y. N., Chen, Z., Deng, Z., Firestone, B., … LaMarche, M. J. (2016). Allosteric Inhibition of SHP2: Identification of a Potent, Selective, and Orally Efficacious Phosphatase Inhibitor. J Med Chem, 59(17), 7773–7782. doi:10.1021/acs.jmedchem.6b00680

Gavrieli, M., Watanabe, N., Loftin, S. K., Murphy, T. L., & Murphy, K. M. (2003). Characterization of phosphotyrosine binding motifs in the cytoplasmic domain of B and T lymphocyte attenuator required for association with protein tyrosine phosphatases SHP-1 and SHP-2. Biochem Biophys Res Commun, 312(4), 1236–1243.

Hartman, Z. R., Schaller, M. D., & Agazie, Y. M. (2013). The tyrosine phosphatase SHP2 regulates focal adhesion kinase to promote EGF-induced lamellipodia persistence and cell migration. Mol Cancer Res, 11(6), 651–664. doi:10.1158/1541-7786.MCR-12-0578

Hof, P., Pluskey, S., Dhe-Paganon, S., Eck, M. J., & Shoelson, S. E. (1998). Crystal structure of the tyrosine phosphatase SHP-2. Cell, 92(4), 441–450. doi:S0092-8674(00)80938-1 [pii]

Keegan, K., & Cooper, J. A. (1996). Use of the two hybrid system to detect the association of the protein-tyrosine-phosphatase, SHPTP2, with another SH2-containing protein, Grb7. Oncogene, 12(7), 1537–1544.

Klinghoffer, R. A., & Kazlauskas, A. (1995). Identification of a putative Syp substrate, the PDGF beta receptor. J Biol Chem, 270(38), 22208–22217.

Kong, M., Mounier, C., Dumas, V., & Posner, B. I. (2003). Epidermal growth factor-induced DNA synthesis. Key role for Src phosphorylation of the docking protein Gab2. J Biol Chem, 278(8), 5837–5844. doi:10.1074/jbc.M208286200

Kusano, K., Thomas, T. N., & Fujiwara, K. (2008). Phosphorylation and localization of protein-zero related (PZR) in cultured endothelial cells. Endothelium, 15(3), 127–136. doi:10.1080/10623320802125250

LaRochelle, J. R., Fodor, M., Vemulapalli, V., Mohseni, M., Wang, P., Stams, T., … Blacklow, S. C. (2018). Structural reorganization of SHP2 by oncogenic mutations and implications for oncoprotein resistance to allosteric inhibition. Nat Commun, 9(1), 4508. doi:10.1038/s41467-018-06823-9

LaRochelle, J. R., Fodor, M., Xu, X., Durzynska, I., Fan, L., Stams, T., … Blacklow, S. C. (2016). Structural and Functional Consequences of Three Cancer-Associated Mutations of the Oncogenic Phosphatase SHP2. Biochemistry, 55(15), 2269–2277. doi:10.1021/acs.biochem.5b01287

Li, S., Couvillon, A. D., Brasher, B. B., & Van Etten, R. A. (2001). Tyrosine phosphorylation of Grb2 by Bcr/Abl and epidermal growth factor receptor: a novel regulatory mechanism for tyrosine kinase signaling. EMBO J, 20(23), 6793–6804. doi:10.1093/emboj/20.23.6793

McAlister, G. C., Nusinow, D. P., Jedrychowski, M. P., Wuhr, M., Huttlin, E. L., Erickson, B. K., … Gygi, S. P. (2014). MultiNotch MS3 enables accurate, sensitive, and multiplexed detection of differential expression across cancer cell line proteomes. Anal Chem, 86(14), 7150–7158. doi:10.1021/ac502040v

Meyers, R. M., Bryan, J. G., McFarland, J. M., Weir, B. A., Sizemore, A. E., Xu, H., … Tsherniak, A. (2017). Computational correction of copy number effect improves specificity of CRISPR-Cas9 essentiality screens in cancer cells. Nat Genet, 49(12), 1779–1784. doi:10.1038/ng.3984

Mootha, V. K., Lindgren, C. M., Eriksson, K. F., Subramanian, A., Sihag, S., Lehar, J., … Groop, L. C. (2003). PGC-1alpha-responsive genes involved in oxidative phosphorylation are coordinately downregulated in human diabetes. Nat Genet, 34(3), 267–273. doi:10.1038/ng1180

Niihori, T., Aoki, Y., Ohashi, H., Kurosawa, K., Kondoh, T., Ishikiriyama, S., … Matsubara, Y. (2005). Functional analysis of PTPN11/SHP-2 mutants identified in Noonan syndrome and childhood leukemia. J Hum Genet, 50(4), 192–202. doi:10.1007/s10038-005-0239-7

Ronnstrand, L., Arvidsson, A. K., Kallin, A., Rorsman, C., Hellman, U., Engstrom, U., … Heldin, C. H. (1999). SHP-2 binds to Tyr763 and Tyr1009 in the PDGF beta-receptor and mediates PDGF-induced activation of the Ras/MAP kinase pathway and chemotaxis. Oncogene, 18(25), 3696–3702. doi:10.1038/sj.onc.1202705

Roskoski, R., Jr. (2005). Src kinase regulation by phosphorylation and dephosphorylation. Biochem Biophys Res Commun, 331(1), 1–14. doi:10.1016/j.bbrc.2005.03.012

Saxton, T. M., Henkemeyer, M., Gasca, S., Shen, R., Rossi, D. J., Shalaby, F., … Pawson, T. (1997). Abnormal mesoderm patterning in mouse embryos mutant for the SH2 tyrosine phosphatase Shp-2. EMBO J, 16(9), 2352–2364. doi:10.1093/emboj/16.9.2352

Subramanian, A., Tamayo, P., Mootha, V. K., Mukherjee, S., Ebert, B. L., Gillette, M. A., … Mesirov, J. P. (2005). Gene set enrichment analysis: a knowledge-based approach for interpreting genome-wide expression profiles. Proc Natl Acad Sci U S A, 102(43), 15545–15550. doi:10.1073/pnas.0506580102

Tang, T. L., Freeman, R. M., Jr., O’Reilly, A. M., Neel, B. G., & Sokol, S. Y. (1995). The SH2-containing protein-tyrosine phosphatase SH-PTP2 is required upstream of MAP kinase for early Xenopus development. Cell, 80(3), 473–483.

Tartaglia, M., & Gelb, B. D. (2005). Germ-line and somatic PTPN11 mutations in human disease. Eur J Med Genet, 48(2), 81–96. doi:10.1016/j.ejmg.2005.03.001

Xu, D., & Qu, C. K. (2008). Protein tyrosine phosphatases in the JAK/STAT pathway. Front Biosci, 13, 4925–4932.

Yokosuka, T., Takamatsu, M., Kobayashi-Imanishi, W., Hashimoto-Tane, A., Azuma, M., & Saito, T. (2012). Programmed cell death 1 forms negative costimulatory microclusters that directly inhibit T cell receptor signaling by recruiting phosphatase SHP2. J Exp Med, 209(6), 1201–1217. doi:10.1084/jem.20112741

Zhao, R., Guerrah, A., Tang, H., & Zhao, Z. J. (2002). Cell surface glycoprotein PZR is a major mediator of concanavalin A-induced cell signaling. J Biol Chem, 277(10), 7882–7888. doi:10.1074/jbc.M111914200

Zhao, R., & Zhao, Z. J. (2000). Dissecting the interaction of SHP-2 with PZR, an immunoglobulin family protein containing immunoreceptor tyrosine-based inhibitory motifs. J Biol Chem, 275(8), 5453–5459.

